# Unifying N- and O-glycopeptide dissociation methods with expedient electron transfer dissociation

**DOI:** 10.64898/2026.10.12.758703

**Authors:** Haley M. Schramm, Tim S. Veth, Joshua D. Hinkle, Jingjing Huang, David Bergen, Rafael D. Melani, Graeme C. McAlister, Christopher Mullen, Nicholas M. Riley

## Abstract

The glycoproteome is highly heterogeneous, where each glycosite can be modified by potentially hundreds of glycan compositions. This complexity necessitates methods that can provide glycosite-specific characterization in a high-throughput manner. Electron transfer dissociation (ETD) is particularly valuable for glycopeptide analysis, where it is critical for glycosite localization in O-glycopeptides and often produces higher quality spectra than higher energy collisional dissociation (HCD) for N-glycopeptides. Despite this, lower quality HCD spectra are often favored in high-throughput glycoproteomics experiments because the near instantaneous fragmentation of HCD results in faster MS/MS spectral acquisition relative to ETD. In short, overhead times for ETD reactions limit acquisition rates and sensitivity that can outweigh its advantages. Here we implement changes to ion handling, scan parallelization, and ion-ion reaction conditions to increase ETD MS/MS acquisition rates on the Orbitrap Tribrid MS platform. We explore fundamental parameters of ETD ion-ion reactions, including reaction q-values (i.e., Mathieu q-values) and ion-ion reaction time, showing that we can reduce the overhead of ion-ion reaction scan functions and improve the overall synchronization of data acquisition. These optimizations lead to faster spectral acquisition times for ETD MS/MS spectra while maintaining high spectral quality. Ultimately, these improvements allow the ETD scan functions to be nested within ion accumulation times needed per MS/MS spectrum, enabling ETD spectral acquisition to be parallelized within the duty cycle of the instrument and removing the major time cost to doing ETD-based fragmentation. We argue that this expedient implementation of ETD makes ETD-based fragmentation a universally beneficial dissociation method for both N- and O-glycopeptides.

## INTRODUCTION

Electron transfer dissociation (ETD) is a tandem mass spectrometry (MS/MS) method that involves reacting multiply charged precursor cations with singly charged radical reagent anions, typically within the confines of an rf-trapping device.^1–4^ Transfer of an electron from the reagent anion to the precursor cation creates an odd-electron species that undergoes free-radical-driven cleavage of N–C_α_ bonds along peptide backbone of precursor ions. This gas-phase dissociation generates even electron c-type and odd electron z^•^-type product ions that provide peptide sequence information. Importantly, the radical dissociation of peptide backbone bonds in ETD largely retains labile modifications intact at modified amino acid residues, making it a valuable MS/MS method for analyses of proteins and their post-translational modifications (PTMs).^5–14^

ETD has emerged as an indispensable tool for glycoproteomics in particular.^15–18^ Protein glycosylation comprises a complex family of PTMs that includes multiple classes (e.g., N- and O-glycosylation) and hundreds of possible glycan compositions that modify glycosites in a largely non-templated fashion. This heterogeneity is an inherent feature of glycosylation that acts as an important modulator of biological function.^19–21^ It also introduces numerous technical challenges that make glycoproteome characterization significantly more demanding than other proteome-centric analyses. The extensive microheterogeneity of glycosylation, i.e., the diversity of glycans that occupy any given site, necessitates intact glycopeptide analysis where glycan modifications can be localized to a specific amino acid(s) within a peptide sequence.^22–24^ Even so, most intact glycopeptide analyses do not fully capture the topology of glycans themselves, leaving structural characterization of glycopeptides as an elusive goal for most high-throughput glycoproteomic experiments.^25^ Our work here does not address this limitation in current glycoproteomic methodology but instead focuses on using tandem MS to assign site-specific glycan compositions, as is common in most intact glycopeptide analyses. Glycosidic linkages are more labile than peptide bonds, meaning traditional collision-based methods like beam-type collisional dissociation (sometimes called higher energy collisional dissociation, HCD) can be limited in their ability to provide site-specific information. ETD, on the other hand, has excelled at this task for both N- and O-glycopeptides, especially when combined with supplemental activation strategies that improve ETD dissociation efficiencies.^26–35^ ETD-based methods serve as the *de facto* method for O-glycosite localization and often generate more informative spectra for N-glycopeptides than HCD.^35–40^ Indeed, the impact of electron-driven dissociation for glycopeptides (and for other features of the proteome) has driven numerous innovations to bring a range of electron-activated dissociation (ExD) methods to a wide array of MS instrument platforms.^41–53^

Despite clear advantages, ETD-based methods are underutilized because ion-ion reactions have high scan overhead times that slow down spectral acquisition. The result of this time cost is that most high-throughput glycoproteomics experiments favor HCD over ETD-based methods because significantly more HCD MS/MS spectra are often sufficient to translate to more identifications, regardless of quality.^25,35,54,55^ Other methods use combinations of collision-based and electron-based methods to minimize detrimental acquisition rates associated with ETD scans while still collecting ETD spectra for a subset of precursor ions.^56–62^ While valuable, these methods “double up” by taking multiple MS/MS spectra per precursor ion, limiting sensitivity. Thus, improving ETD-based methods to mitigate these limitations would be advantageous for glycoproteomics.

ETD spectral acquisition is largely dictated by both the reaction time required to generate sequence-informative product ions and the overhead steps associated with transmitting and storing ions of different polarities. Here, our goal was to investigate how we could expedite ETD spectral acquisition to benefit glycoproteome characterization through alteration of the parameters governing the ion-ion reaction itself and the scan acquisition sequences leading up to the ETD reaction. We show how shortening reaction times based on kinetic considerations can generate sufficient product ion signal-to-noise with faster spectral acquisition rates, and we investigate how balances in parallelization, ion accumulation time, and ion-ion reaction time can be leveraged to improve glycoproteome sampling with ETD-based methods. Ultimately, our expedient ETD-based methods show that ETD reactions can be conducted with overhead times lower than precursor ion accumulation times, enabling parallelization of ETD scan functions and significantly reducing the time costs of performing ETD-based fragmentation. This improvement makes previously observed benefits of HCD fragmentation less dominant and positions ETD-based fragmentation as an ideal dissociation method for both N- and O-glycopeptides.

## EXPERIMENTAL PROCEDURES

### Sample Preparation

#### Glycocarrier

The O-glycocarrier^63^ was resuspended, reduced, and alkylated in lysis buffer [1% sodium deoxycholate, 10 mM tris(2-carboxyethyl)phosphine hydrochloride, 40 mM chloroacetamide, and 100 mM Tris, pH 8.5 supplemented with protease inhibitor (cOmplete mini EDTA-free, Roche)]. Proteolytic digestion was completed overnight at 37 °C with using sequencing trypsin (Promega) in a 1:50 enzyme:protein ratio by weight. Following digestion, SDC was precipitated twice with 2% formic acid (FA), after which samples were desalted using Strata-X cartridges (Phenomenex) following the vendor-supplied protocol. The resulting sample was dried by vacuum centrifugation and stored at −70 °C until mass spectrometry analysis

#### Mucin mix O-glycoproteomics

Recombinant mucin glycoproteins were sourced from AcroBio systems. Ten micrograms of each leukosialin (CD43) [P16150], mucin-16 [Q8WXI7], P-selectin glycoprotein ligand 1 [Q14242], podocalyxin [O00592], and platelet glycoprotein1b alpha chain (GP1bα) [P07359] were pooled together and digested with the O-glycoprotease IMPa (New England Biolabs) for 3 hours. The sample was then reduced with tris(2-carboxyethyl)phosphine (TCEP) and carbamidomethylated with chloroacetamide (CAA) prior to digesting with trypsin overnight. The resulting glycopeptide mixture was desalted via SPE (Phenomenex Strata-X cartridges) as described previously, dried by vacuum centrifugation, and stored at −70 °C until mass spectrometry analysis.^64^ The sample was reconstituted in 0.1% formic acid and ∼300 ng of starting material was injected on column.

#### Serum N-glycoproteomics

Human serum (Invitrogen, catalog #31876) was reconstituted in 1% sodium deoxycholate (SDC), 10 mM tris(2-carboxyethyl)phosphine hydrochloride, 40 mM chloroacetamide, and 100 mM Tris, pH 8.5, supplemented with protease inhibitor (cOmplete mini EDTA-free, Roche) to reduce and alkylate the sample. The protein samples were digested overnight at 37°C using trypsin at a1:50(w/w) trypsin/protein ratio, desalted, and glycoenriched with mixed-mode anion exchange cartridges as described previously.^62^ The resulting sample was dried by vacuum centrifugation and stored at −70 °C until mass spectrometry analysis. The sample was reconstituted in 0.1% formic acid and ∼2.5 ug of starting material was injected on column.

### Data Acquisition

#### Direct infusion of the glycocarrier, Figure 1

The digested glycocarrier was directly infused by static spray using a borosilicate nanoelectrospray capillary pulled in house. The voltage applied to the needle was between 0.8 and 1.5 kV. MS2 spectra were acquired via a custom diagnostic. This function iterated through 10 scans at varying reaction times with user defined q-values and tau values and were saved to a raw file and output into a .txt file. This data was acquired on an Orbitrap Ascend Tribrid MS modified to allow adjustment of ion-ion reaction q-values.

**Figure 1.**
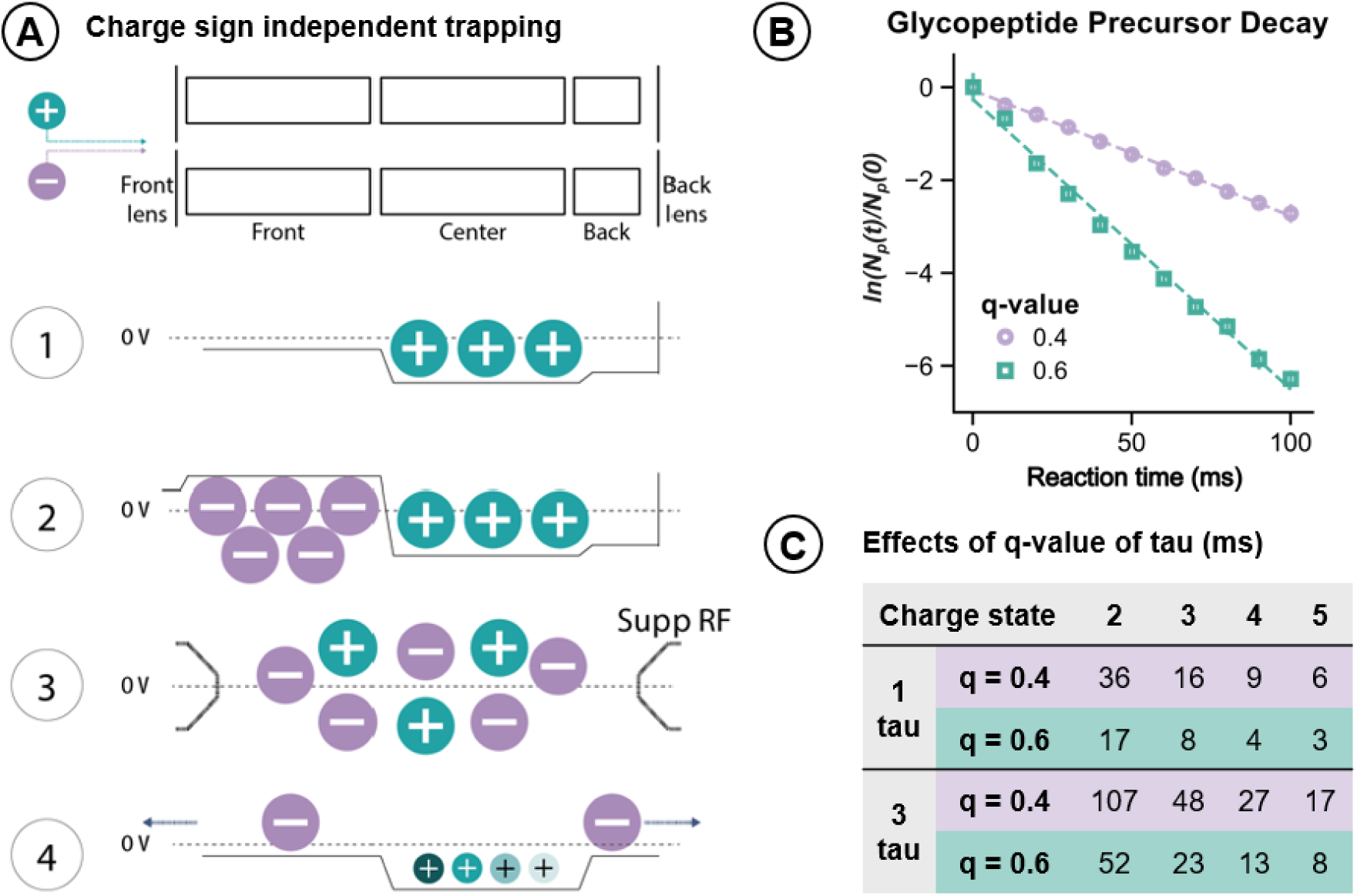
**Effects of ion-ion reaction parameters**. **A)** The ETD ion-ion reaction process occurs in the high-pressure linear ion trap and includes (1) accumulating precursor cations in the center section, (2) accumulating reagent anions in the front section, (3) charge sign independent trapping (CSIT) using a supplemental RF applied to the end lenses to mix the ion populations, and (4) quenching the reaction to retain positively charged product ions. The ETD reaction time is the duration of CSIT in step three, which is where the trapping potential, and thus q-value, used can be changed to alter the degree of ion cloud overlap. **B)** Raising the q-value used during the ETD reaction from 0.4 to 0.6 depletes the precursor ion population at a faster rate, where the slope is -k, i.e., the negative value of the rate constant. Data shown here is the doubly charged cation of the glycopeptide AEAAAT[HexNAc]PAPAK, m/z 600.81. C) Tau values (τ), the inverse of the rate constant k, provide ETD reaction time constants to standardize precursor depletion, and thus, extent of ETD reactions. The table here provides multiples of tau, 1τ and 3τ, for precursor ion charge states z = 2-5 for reaction conditions using a q-value of 0.4 and 0.6 relative to the reagent anion at m/z 202.

#### ETD evaluations in Figure 2 and 3a

The mucin O-glycoproteomics sample was analyzed on a Thermo Scientific Orbitrap Ascend Tribrid Mass Spectrometer (Waltham, MA) coupled to a Thermo Fisher Scientific Vanquish Neo UHPLC. Samples were reconstituted in 0.1% formic acid and loaded onto a C18 trap column using 100% mobile phase A (0.1% formic acid). A 55-minute gradient ramped until 28% mobile phase B (99.9% ACN with 0.1% FA) on a C18 analytical column (75 μm ID x 25 cm, IonOpticks) followed by a 5-minute column wash with 99% mobile phase B and equilibrated with mobile phase A. The sample was ionized using a Nanospray Flex ionization source (Thermo Fisher Scientific) held at +2.0 kV compared to ground, and the inlet capillary temperature was held at 275 °C. Full MS spectra were acquired in the Orbitrap with a resolving power of 60,000 or 120,000 at m/z 200 and an m/z range of 600-1600 (by quadrupolar isolation). The full scan AGC target was set to standard (100%, 400,000 charges) with a maximum injection time of 128 or 251 ms. Precursor ions with charge states z = 2-5 and an MS1 intensity greater than 25,000 or 50,000 were selected with the quadrupole (0.7 or 1.6 m/z isolation windows) for MS/MS in data-dependent fashion using a dynamic exclusion duration of 45 or 60 seconds. MS/MS spectra were acquired with varying fragmentation conditions (e.g., ion-ion reaction q-values, reaction times, supplemental collision energies) as defined in discussion of results. MS/MS methods utilized Orbitrap mass analysis with a 200% AGC target (100,000 charges), a resolving power of 30,000 or 60,000 at m/z 200, and a maximum injection time of 64 or 123 ms. All ETD reagent anion AGC target values were set to 500,000 charges, and reagent anion AGC targets were always met prior to an ETD reaction. sceHCD spectra were collected with an m/z 120-2000 range, while ET(hc)D spectra were collected with a m/z 120-3600 range based on previous analyses that show the benefit of higher mass ranges for ETD-based spectra to measure glycan-retaining product ions.^64^ Control of ion-ion reaction parameters was enabled within the method by an instrument patch that allowed for the scan description to define the time constants and q-value to use for a given scan event.

**Figure 2.**
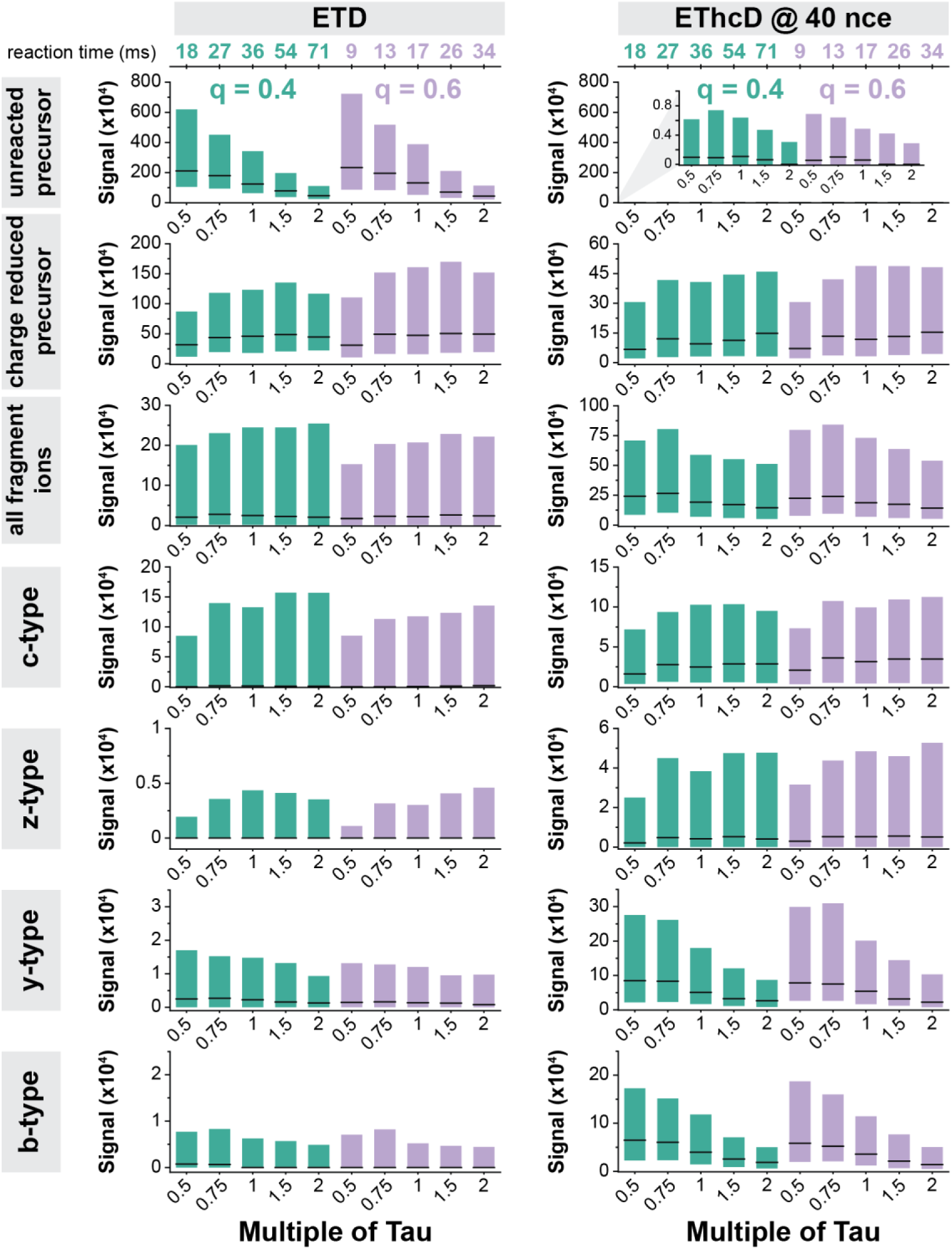
**Effects of ETD reaction parameters on product ion types for doubly charged O-glycopeptide precursor ions**. Box plots show median (black horizontal line) and quartile values (boxes) for various ion types for ETD (left) and EThcD with a supplemental activation of 40nce (right) for doubly charged O-glycopeptides at various reactions conditions. Each column of graphs shows data for ETD or EThcD, while each row shows data for specific ion types, as shown with gray label boxes above the columns and to the left of the rows. Ion types shown in order from bottom to top are unreacted precursor, charge reduced precursor, all fragment ion types (c, z, y, b) summed, and each fragment ion type separately. Within each set of box plots, reaction conditions using either a q-value of 0.4 (green) or 0.6 (purple) are shown for different multiples of τ, with the corresponding reaction time in ms given at the top of each column of graphs. Analyses represent an average of ∼475 O-glycopeptides. All y-axes are scaled to show signal x 104 but the y-axes are scaled differently as appropriate for signal from each ion type. The graph for the unreacted precursor in EThcD is scaled the same as its analogous ETD plot, while the inset shows a zoom in to enable more straightforward visualization. Each of these MS/MS methods used a precursor ion AGC target of 100,000 charges, a reagent AGC target of 500,000 charges, and Orbitrap mass analysis with 60,000 resolving power, and a 123 ms precursor ion maximum injection time.

**Figure 3.**
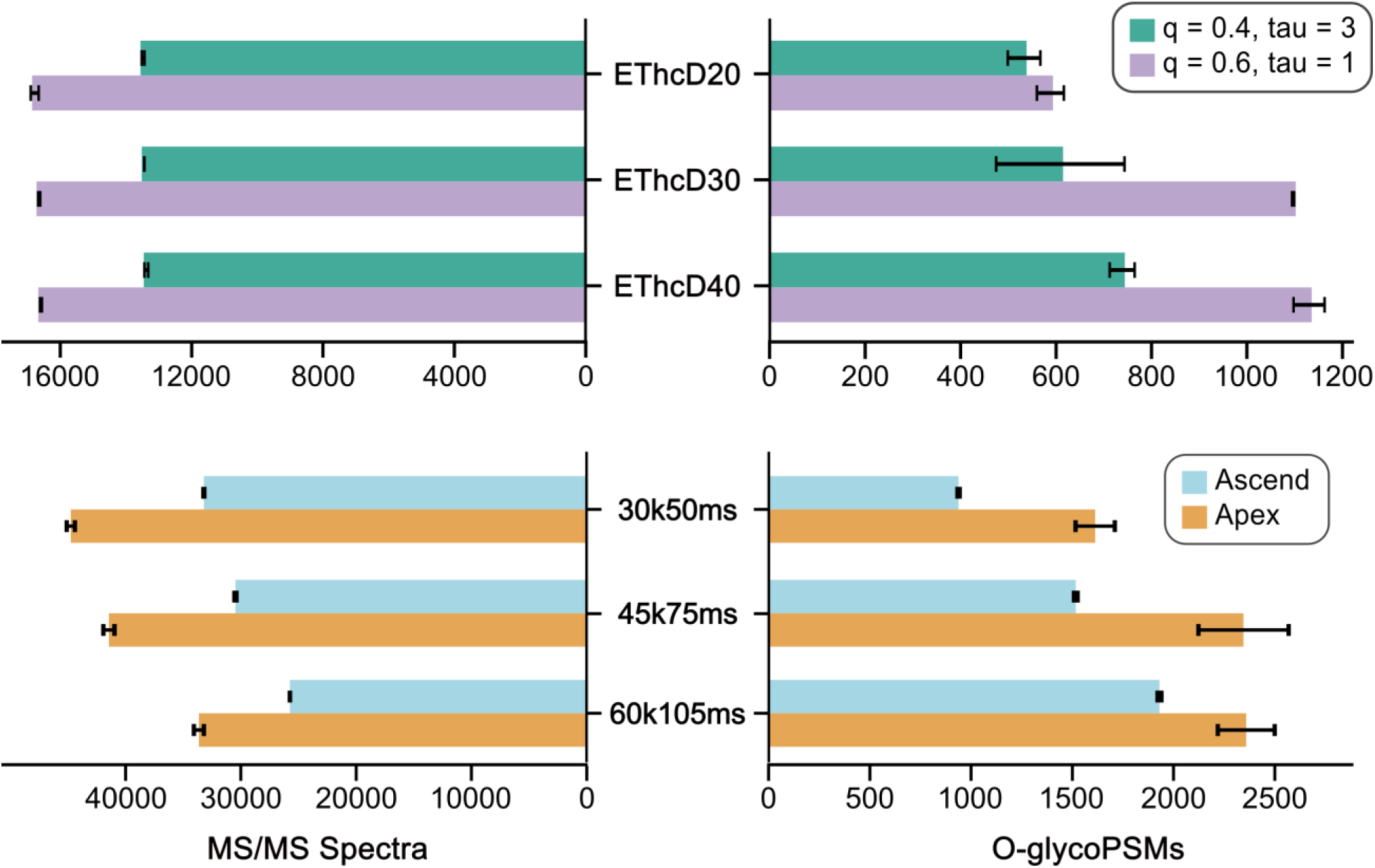
Translating new ETD and ion pipelining parameters to glycoproteomics. **A)** Counts of MS/MS spectra and O-glycopeptide spectral matches (O-glycoPSMs) are shown for DDA methods using EThcD for MS/MS fragmentation with either 20, 30, or 40 nce as the supplement activation to analyze a complex mixture of O-glycopeptides. Green bars show data when a q-value of 0.4 and 3τ were used for the ETD reaction (the standard on the Orbitrap Ascend), while purple bars show data when a q-value of 0.6 was used for the ETD reaction. Each of these MS/MS methods used an AGC target of 100,000 charges and Orbitrap mass analysis with 30,000 resolving power and 64 ms maximum injection time. **B)** Counts of MS/MS spectra and O-glycoPSMs are shown for DDA methods using EThcD for MS/MS fragmentation with 40 nce as the supplement activation. Methods varied in their resolving power and their total precursor ion accumulation time that was scaled with transient length. Blue bars show data collected with the standard ion pipelining and ETD reaction parameters on the Orbitrap Ascend, while orange bars show data with the updated ETD reaction parameters and ion pipelining on the Orbitrap Apex. All data were collected in duplicate, and bars show the average of the two replicates, whose values are indicated by the error bars.

#### Ion pipelining evaluation and further Orbitrap Apex testing, Figures 3b, 4, and 5

Data comparing the improved ion pipelining approaches were obtained using single modified Tribrid MS system that was alternated between operating as an Oribtrap Ascend or utilizing prototypical configuration equivalent to an Orbitrap Tribrid Apex MultiOmics MS (with the modifications more explicitly discussed elsewhere^65^). All user-defined method parameters between the Ascend versus Apex modes of operation remained constant between methods. LC separations used 65-minute gradient ramped until 23% mobile phase B (80% ACN with 20% 0.1% FA) on a C18 analytical column (75 μm ID x 50 cm, IonOpticks) followed by a second ramp to 40% B for 20-minutes and then a 12-minute column wash with 95% mobile phase B and equilibrated with mobile phase A. The sample was ionized using a Nanospray Flex ionization source (Thermo Fisher Scientific) held between +1.8 and 2.2 kV compared to ground, and the inlet capillary temperature was held at 275 °C. MS and MS/MS parameters remained the same as in previous sections, but the resolving power and maximum injection times were varied as noted in discussion of Figure 3. Experiments for Figures 4 and 5 used 45,000 resolving power for MS/MS Orbitrap mass analysis with 75 ms maximum ion injection time.

**Figure 4.**
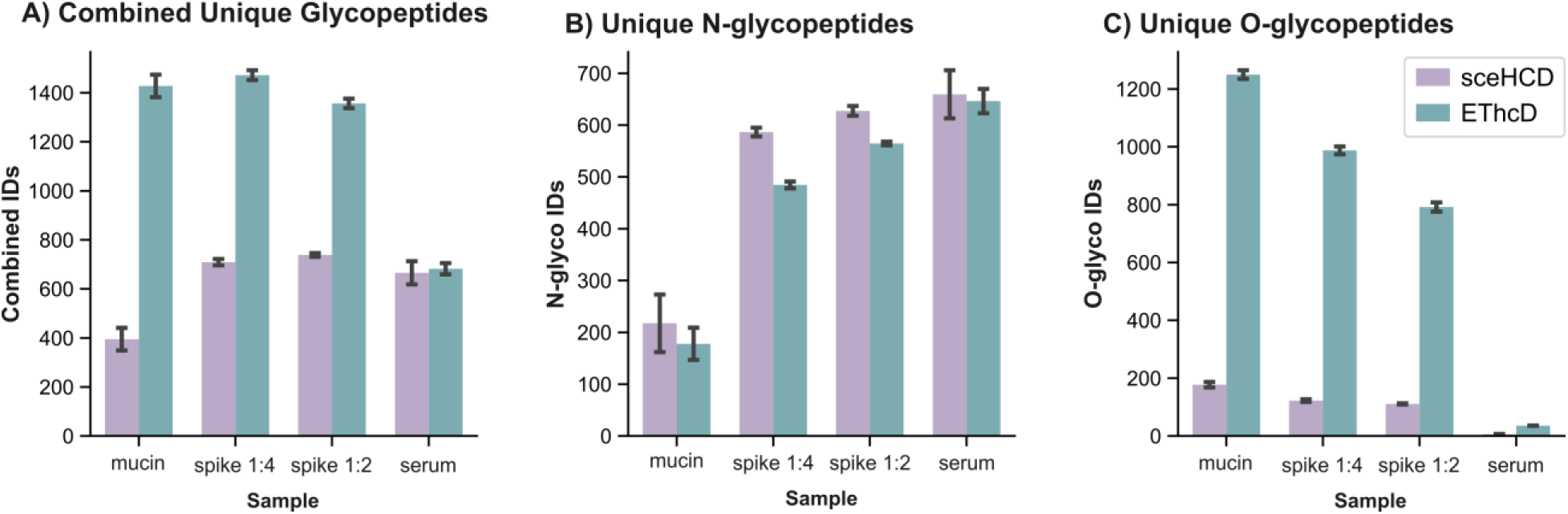
Comparing sceHCD and EThcD methods for N- and O-glycoproteomics on the Orbitrap Apex. Bar graphs showing the number of **A)** combined glycopeptide identifications, **B)** N-glycopeptide identifications, and **C)** O-glycopeptide identifications from DDA methods on the Orbitrap Apex using either sceHCD or EThcD40 for MS/MS dissociation. The samples analyzed were either glycopeptides enriched from human serum, a digest of recombinant mucin glycoproteins, or a combination of both to create a complex mixture of both N- and O-glycopeptides. The sceHCD results are shown in purple and EThcD in green. Data were collected in duplicate, and bars show the average of the two replicates, whose values are indicated by the error bars.

**Figure 5.**
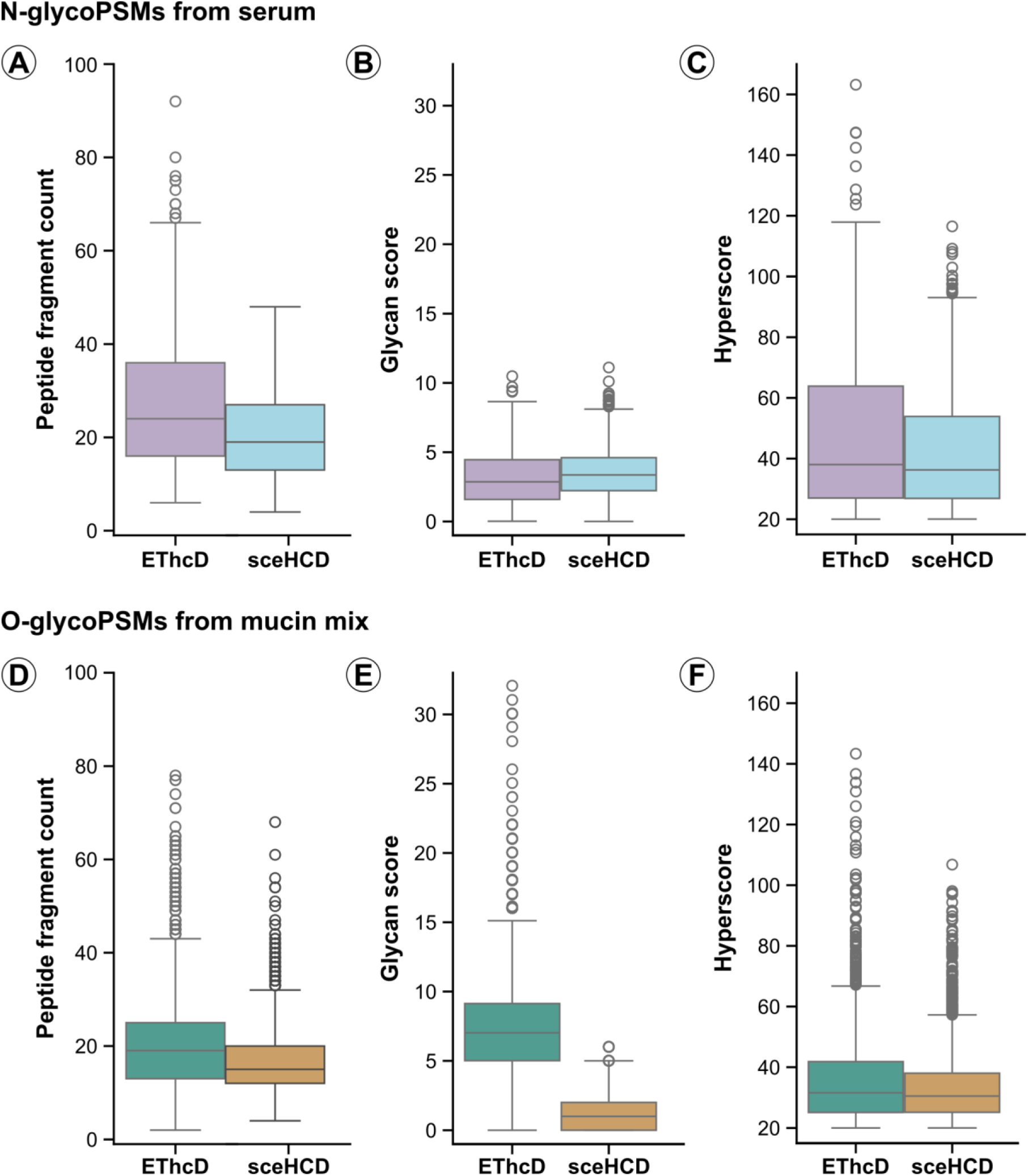
Comparing spectral quality for EThcD and sceHCD N- and O-glycopeptide spectra. Top panels A, B, and C compare fragment ion counts, MSFragger glycan scores, and MSFragger Hyperscores, respectively, for EThcD and sceHCD MS/MS spectra for serum N-glycoPSMs. Bottom panels show the same three comparisons for EThcD and sceHCD MS/MS spectra for mucin O-glycoPSMs. Peptide fragment ion counts include c/z- and b/y-type fragment ions for EThcD and b/y-type fragments for sceHCD. Box plots show quartile ranges, with whiskers showing data points within 1.5 times the interquartile range and outliers shown with open circles. The axes for each of these comparisons are kept consistent between N-glycopeptides (top) and O-glycopeptides (bottom). Note that the y-axes for Hyperscore distributions do not include a zero value.

### Data Analysis

All searches were performed on a PC running Windows 10, with two 2.20 GHz Intel Xeon Silver 4114 CPU processors with 64 Gb of installed RAM, or on a PC running Windows 10, with two 3.00 GHz Intel Xeon W-2295 CPU processors with 128 Gb of installed RAM. Sixteen cores were used per search. O-glycopeptide raw data were searched using O-Pair Search implemented in MetaMorpheus, which is available at https://github.com/smith-chem-wisc/MetaMorpheus.^66^ The spectra were searched against a FASTA file containing Uniprot-derived sequences from all five proteins as described by their sequences from the vendor, which was concatenated with decoy peptides by reversing sequences of the forward database. The “O-Glycopeptide Search” option was selected, where the O-glycopeptide search feature was enabled with an O-glycan database of 22 common O-glycans. The “Keep top N candidates” was set to 10, and the Data Type was set to EThcD or ETD. A mass tolerance of ±10 ppm was used for precursor ions and ±20 ppm for product ions. Methionine oxidation was specified as a variable modification with a maximum of 2 variable modifications per peptide, while carbamidomethylation of cysteine was set as a fixed modification. A combined IMPa and trypsin specificity was defined as previously described,^67^ up to 12 missed cleavages were used (due to IMPa’s cleavage behavior), and the maximum modification isoforms permitted was 1024. The oglyco.psmtsv results file was used, and the data was further filtered using C# scripts^35^, some of which used the C# Mass Spectrometry Library (CSMSL, https://github.com/-dbaileychess/CSMSL). Only the highest quality (Level 1, localized) identifications were used for further analysis. N-glycopeptide identifications were made by searching using MSFragger-Glyco^68^. All spectra were searched using the default glyco-N-hybrid workflow. Both workflows used a mass tolerance of 20 ppm, isotope error of 0-2, strict trypsin enzyme with a maximum of 2 missed cleavages, peptide length of 7-50 amino acids, and 400-5000 Da mass range. Carbamidomethylation on cysteine was set as a fixed modification and oxidation on methionine was set as a variable modification, with a maximum of 3 variable modifications per peptide. A fasta database of the human proteome was downloaded from UniProt (20456 protein entries), and a database of 253 human N-glycans was used from the provided MSFragger-Glyco glycan databases. The glyco-N-HCD workflow searched for b, y, and Y ions. The glyco-N-hybrid workflow searched for b, y, c, z, and Y ions. Results were filtered for a 1% peptide false discovery rate (q-value). Oxonium ion intensities were extracted using GlyCounter with standard settings.^69^ The mass spectrometry raw data and search results have been deposited to the ProteomeXchange Consortium via the MassIVE partner repository with the dataset identifier MSV000102759.

## RESULTS AND DISCUSSION

### Expedient electron transfer dissociation by altering ion-ion reaction kinetics

Ion-ion reactions for ETD MS/MS spectra are conducted in the linear ion trap on Orbitrap Tribrid MS systems, where charge sign independent trapping is conducted with a supplemental RF on the end lenses to mix the ion populations (**Figure 1a**). The rate equation for a second order ion-ion reaction where reactant precursor cations (P) plus reagent anions (R) react to form products, can be written as shown in Equation 1:

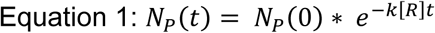

where Np(0) represents the initial precursor population, Np(t) is the amount of precursor remaining at time t, [R] is the number density of the reagent ion cloud, and k is the ion-ion capture rate coefficient. As described by McLuckey, Stephenson, and co-workers^70–73^, the ion-ion capture rate coefficient is defined by Equation 2:

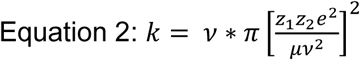

where ν is the relative velocity of the oppositely charged ions, µ is the reduced mass of the collision partners, and z_1_ and z_2_ are the charges of the reaction partners, and e is the elementary charge, and has been demonstrated to take on a charge-state squared relationship.

It is typical to recast the reaction rate coefficient into a pseudo-first order form for measurement purposes, according to Equation 3, by operating under reaction conditions where the reagent anions are in a large excess to the reaction precursor cations, [R] >> [P].

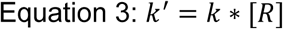

Prior work from Rose et al. sought to improve ETD functionality through a calibration routine that scaled reaction times in a charge-state dependent manner based on time constants (denoted as tau, τ, where τ = 1/k’).^74^ These time constants are the reciprocal of the charge state dependent ETD reaction rate constant k’. The result of that work was the observation of maximization of product ion signal at approximately two-to-three time constants (2-3τ), although acceptable fragment ion signal-to-noise and identifications can still be achieved with lower reaction times (**Figure S1**). That investigation set the foundation for calibrated ETD reaction times on all commercial Orbitrap Tribrid MS systems to use 3τ when calibrating charge-dependent ETD settings. This combination of parameters, however, can lead to longer reaction times than required for identification, as shown by our recent studies that optimized scan times on the dual ion routing multipole-geometry of the Orbitrap Ascend.^52^

As can be seen from Equation 2 and Equation 3, the pseudo-first order reaction rate coefficient is a product of the reagent anion density and the relative velocity of the reacting ion pairs. An increase in k’ can be realized by increasing the reagent anion density and/or decreasing the relative velocity of the reacting ion pairs, and both parameters are linked to the Mathieu q-value during the ion-ion reaction.

The vast majority of ion-ion reactions, including those on commercial Orbitrap Tribrid MS systems up until now, are conducted with a q-value at 0.4 relative to the fluoranthene reagent anion’s m/z 202 value, providing a low mass cutoff at ∼ m/z 90 that is suitable for retaining most useful peptide-derived product ions in proteomics.^75,76^ As shown in **Figure 1b**, as the reaction q-value is increased, we observe an increase in the reaction kinetics up to a Mathieu q-value of 0.6. While it is difficult to estimate the pseudo-potential well depth in an ion trapping device at q-values above about 0.4, it is understood that the well depth should increase monotonically over the range of q-values investigated, and that the increased well depth provides higher radial confinement, i.e. density of the reagent anion cloud. Since kinetics are observed to increase over this q value range, we postulate that the rate of change of the reagent density is changing faster than any increase in the relative velocity (ν^3^) term of Equation 2 that may be anticipated, leading to a net increase in the reaction rate. Similar trends in kinetics have been previously observed by Compton et al. at increased ion-ion reaction q values.^73^

We used a doubly charged O-glycopeptide precursor ion derived from the engineered O-glycocarrier to show this effect for glycopeptides that are interest for this study (**Figure 1b**).^63^ **Figure S2** provides similar data for other glycoforms of this peptide to show how this trend holds across different glycopeptide species, and this behavior applies to precursor ions regardless of biomolecule type (**Figure S3**).^77^ The slope of the depletion curve provides the ETD reaction rate constant k^’^, so the steeper the slope, the faster the reaction. Conducting ETD reactions with a q-value of 0.6 generates a steeper slope than reactions with a q-value of 0.4, showing that the reaction proceeds at a faster rate with a higher q-value, with the reaction rate increasing in a linear fashion over the range of q values studied, (**Figure S4**). The charge state dependent reaction times are calculated from the measurement of k’ at a fixed q-value and precursor charge state (0.4 and 2, respectively, for example) and converted to the charge state and number of precursor depletion time constants through Equation 4:

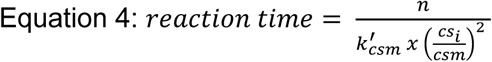

where n is the number of multiples of τ, k’_csm_ is the pseudo first order rate coefficient for the charge state measured, and cs_i_ is the charge state in question. **Figure S5** provides further evidence that reaction rates scale with charge state as expected with both the 0.4 and 0.6 q-values used. **Figure 1c** shows the 2-fold reduction in reaction time that can be realized by operating at a higher ion-ion reaction q-value and for fewer time constants (i.e., fewer multiples of τ).

Next, we evaluated how multiples of the time constant (τ), i.e., different ETD reaction times, affect signal for several ion types for glycopeptide precursors (**Figure 2**). These experiments used a back-to-back scan approach we refer to as a “stacked scan” acquisition sequence for MS/MS analysis. These stacked scan sequences were used in LC-MS/MS methods analyzing complex mixtures of O-glycopeptides derived from sequential digests of recombinant mucin glycoproteins with the O-glycoprotease IMPa and trypsin.^67,78^ The stacked scan sequence employed meant that six MS/MS scans were collected for every precursor selected in data-dependent fashion, where each MS/MS scan had a different multiple of τ (0.5, 0.75, 1.0, 1.5, and 2.0, with corresponding reaction times in ms shown at the top of **Figure 2**). These spectra were acquired in randomized order to minimize systematic effects from ion abundance differences that could result during elution from the LC column during acquisition. The sixth MS/MS scan was an HCD spectrum to aid with identification. Each LC-MS/MS analysis used a single q-value (0.4 or 0.6) with various multiples of τ and each injection focused on only a single charge state (z = 2 or z =3). We also performed these experiments using standard ETD and ETD with HCD supplemental activation at 40 nce (EThcD40).

The top row of **Figure 2** shows how intact precursor ion signal is depleted as expected in ETD reactions with higher τ multiples (i.e., longer reaction times). The second row provides evidence for generation and then depletion of charge-reduced precursor ion signal at higher τ multiples, representing the onset of “overreacted” conditions. These relationships demonstrate the complexity of selecting ion-ion reaction times. Sufficient time is needed to generate maximum product ion signal, but the reaction itself consumes charged product ions as well, meaning valuable product ion signal can be unnecessarily depleted at longer reaction times (**Figure S6**), which has also been observed in ECD.^79^ In EThcD40 conditions, the remaining intact precursor population is almost fully fragmented away during the supplemental HCD step. This reflects expected behavior because the intact multiply charged precursor resides at a lower m/z and has more charges, which typically improves HCD fragmentation efficiency. For triply charged O-glycopeptides, we see that intact z = 3 unreacted precursor ions and doubly-charged, singly-charge-reduced precursor ions (z = 2) are depleted by supplemental HCD more than singly-charged, doubly-charge-reduced precursor ions (z = 1) (**Figure S7**). Ultimately, this fragmentation behavior translates to a useful balance between extent of ETD reaction and supplemental activation strategy. Shorter ETD reaction times that generate sufficient c/z-type ion signal but leave unreacted precursor ions that benefit b/y-type product ion and oxonium ion production in EThcD spectra. These oxonium and b/y-type ions are particularly useful in common search strategies for O-glycopeptides.^80,81^ Overall trends showing similar ion signal for reactions using a q-value of 0.4 or 0.6 were observed for triply charge O-glycopeptide precursors (data not shown).

The bottom five rows of Figure 2 compare total fragment ion signal and signal for c-, z-, y-, and b-type fragment ions in ETD and EThcD40 reactions at different multiples of τ. These data show that sufficient fragment ion signal can be generated at lower multiples of τ and that a q-value of 0.6 can provide the same ETD-driven fragmentation quality as longer reaction times at a q-value of 0.4. Standard reaction conditions on the Orbitrap Tribrid MS systems up to this point used a q-value of 0.4 and 3τ (107 ms for z = 2) for reaction time. Figure 2 shows that a q-value of 0.6 and 1τ for reaction time (17 ms for z = 2) can generate roughly equivalent data, representing a ∼6-fold decrease in reaction time needed. Numerous other interesting trends can be observed, including a general leveling off the median c/z-type ion signal after ∼1τ. EThcD can help improve both c/z-and b/y-type ion signal. Converse to c/z-type ions, b/y-type ions decrease as multiples of τ increase. This trend supports the discussion above, where shorter reaction times lead to a larger population of unreacted precursor ions that is converted to b/y-type fragment ions during supplemental HCD. Although the higher q-value does raise the low m/z cutoff (LMCO) for the ETD reaction to ∼135 m/z (**Figure S4)**, the majority of low m/z oxonium ions that are critical in glycopeptide MS/MS spectra^64^ are generated during the supplemental HCD step, not the ETD reaction. After ions are subjected to ETD in the ion trap, they are then shuttled back towards to the C-trap through the back ion routing multipole (IRM). The kinetic energy with which they are shuttled into the back IRM is determined by a DC offset that scales to the supplemental normalized collision energy desired. Thus, the effective LMCO for these collision-generated fragments in glycopeptide spectra remains a function of the q-value relative to the precursor ion m/z used for HCD scan events. For example, the LMCO during the supplemental HCD step that uses a q ≈ 0.1 (within the typical range used for HCD) for the m/z 600.81 precursor ion in Figure 1b would be ∼ m/z 66 when considering Equation 5. This lower q-value relative to the precursor m/z used for supplemental HCD activation effectively retains low m/z oxonium ions in EThcD spectra using an ETD reaction q-value of 0.6 (**Figure S8**).

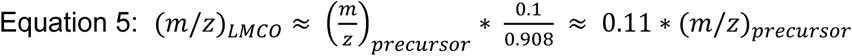

We then extended these observations to analyze the same mixture of O-glycopeptides with traditional DDA methods (**Figure 3a**), i.e., all MS/MS spectra acquired use the same parameters, rather than the stacked scan data acquisition strategy used above. We investigated two ETD reaction conditions, which include current commercial ETD parameters of q = 0.4 and 3τ and our newly optimized parameters of q = 0.6 and 1τ. Based on the balance between remaining precursor and the added value of HCD supplemental activation, we also compared EThcD with 20, 30, and 40 nce. **Figure 3a** shows that the newly optimized ETD reaction parameters (q = 0.6, 1τ) enable collection of significantly more EThcD MS/MS scans regardless of supplemental activation conditions. The faster ETD parameters always enable identification of more O-glycopeptide spectral matches (O-glycoPSMs), but this benefit is far more pronounced when using higher supplemental activation energies for EThcD that convert more of the unreacted precursor to sequence-informative b/y-type ions. Taken together, these data underscore a central thesis of our reaction kinetics experiments: speeding up reaction rates using higher q-values for the ETD reaction and shortening reaction times based on kinetic considerations can generate near-equivalent spectral quality with significantly reduced ETD scan overhead times. Here we demonstrate this for O-glycopeptide precursor ions, which represent one of the more complex use cases, but this trend is expected to hold for most proteomics applications.

### Improved ion pipelining for sensitivity boosts

Although the ion-ion reaction time and kinetic considerations above significantly contribute to overhead times needed to collect ETD spectra, the accumulation of two populations of oppositely charged ion populations and the associated scan functions to execute the ion-ion reaction also require a non-negligible amount of time. The dual IRM architecture of the newest generations of Orbitrap Tribrid MS systems (“Orbitrap Ascend” and “Orbitrap Apex”) helps parallelize some of these functions, but there is still room for improvement for optimizing ion shuttling and scan function parallelization.

Recently implemented advances in ion pipelining developed for the Orbitrap Tribrid Apex improved MS matrix execution efficiency and MS operation synchronization, which increased the portion of each acquisition cycle for available for precursor ion accumulation and reduced the overhead associated with some of the ETD overhead associated with reagent anion injection.^65^ Briefly, the embedded system of the Orbitrap Tribrid MS systems (comprised of a variety of CPUs, FPGAs, and other components) executes organized MS matrices that coordinate MS events, times, and device settings. These components must prepare themselves and align with each other prior to executing an MS matrix. Older generations of Tribrid MS instrument could not perform MS operations in parallel with this overhead time. The Orbitrap Tribrid Apex has a new embedded system that allows ion accumulation to occur in parallel with the preparation time the embedded system requires for setting up the next matrix. This change effectively converts previously unused overhead time into additional parallelizable ion accumulation time on the Orbitrap Tribrid Apex.^65^ Furthermore, the individual steps of injection reagent anions into the linear trap was optimized to reduce the time penalty associated with executing an ETD reaction. Combined with the faster reaction times above, these changes allow for total overhead times for ETD reactions to be fully parallelized within ion accumulation times and mass analysis time needed for Orbitrap MS/MS scans. In other words, these changes nearly eliminate the speed disadvantage that had previously limited broader use of ETD-based fragmentation.

**Figure 3b** summarizes how the ion pipelining changes, in combination with the faster ETD kinetics above, affected EThcD MS/MS spectral acquisition rates and glycopeptide identification. For any given method, the Apex with the faster ETD kinetics and improved ion pipelining generated ∼1.5-fold as many EThcD MS/MS scans than the Ascend. Higher number of MS/MS spectra translated to more O-glycopeptide spectral matches for the Apex methods. As maximum ion accumulation times and concomitant Orbitrap resolving powers increase, both the Ascend and Apex show a decrease in the number of EThcD scan, as expected. This underscores that glycoproteomic methods cannot be driven purely by faster acquisition speeds; they also must balance sensitivity to accumulate enough ions for sufficient signal-to-noise in MS/MS spectra. Increasing ion accumulation time from 50 to 75 to 105 ms on the Ascend generated consistent gains. These same ion accumulation time increases on the Apex leveled out between 75 and 105 ms, showing the complex interplay between methods with fast spectral acquisition rates and sufficient ion accumulation. Using Apex, we are able to acquire on average 10,000 more spectra, which at 45k resolution and 75 ms injection time is 37% more spectra in the same time. Regardless, the faster spectral acquisition times on the Apex consistently enabled more O-glycopeptide identifications with EThcD.

### ETD-based methods as a universal method for glycopeptide identification

With these faster ETD-based methods, we revisited comparisons of stepped collision energy HCD (sceHCD) versus ETD-based methods for glycopeptide identifications. The time costs of ETD-based methods like EThcD are now significantly less dramatic than in previous iterations. **Figure 4** shows how EThcD-only methods with the improved ETD reaction parameters and ion pipeline on the Orbitrap Apex are now competitive with sceHCD methods for N-glycopeptide identifications while also continuing to provide its known advantage for O-glycopeptide identifications. Other metrics from these analyses are available in **Figure S9**. The sample analyzed for this comparison was a glycopeptide enrichment from de-identified pooled human serum combined with O-glycopeptides from an IMPa and trypsin digestion of recombinant mucins, which together provide a reasonable population of N- and O-glycopeptides to evaluate both sceHCD and EThcD methods.

sceHCD and EThcD methods generated comparable numbers of unique N-glycopeptides for the serum-only sample where there are mainly N-glycopeptides present (**Figure 4A**). The similarity in N-glycopeptide identification numbers significantly narrows the gap from previous analyses where sceHCD methods doubled EThcD N-glycopeptide identifications. As the proportion of O-glycopeptides in the sample increased with mucin-derived O-glycopeptides spiked into the serum sample, the EThcD method generated slightly fewer N-glycopeptides identifications (**Figure 4B**). Regardless, the advantage of the EThcD method for O-glycopeptides is clear, with the only localized O-glycopeptide identifications from the sceHCD method coming from O-glycopeptides with a single possible O-glycosite (**Figure 4C**). Ultimately, the EThcD based methods generated more total glycopeptide identifications than sceHCD methods for each serum-mucin combination (**Figure 4A**).

Beyond identification numbers, we investigated the quality of EThcD spectra relative to sceHCD spectra for both N- and O-glycopeptides. We analyzed spectra from the serum-only and mucin-only samples to evaluate the number of fragment ions generated by each method and their search engine score metrics like Glycan score and Hyperscore. **Figure 5** shows these metrics for both fragmentation methods for both N- and O-glycopeptide identifications. EThcD spectra have more annotated ions than sceHCD spectra, which matches trends seen in Figure 2 where four ion types (c/z/y/b-type) are reliably generated by EThcD40 with τ = 1.5 and q = 0.6 conditions used for ETD reactions on the Orbitrap Apex. Glycan scoring differences are primarily observed for O-glycopeptides, and localizing fragments representing a peptide backbone fragment containing the full glycan are exclusively found in EThcD data due to the electron-drive fragmentation of ETD.

## CONCLUSIONS

In this work, we optimized various ion-ion reaction parameters and we examined how these optimized parameters work in concert with updates to ion pipelining on the Tribrid MS system to accelerate the MS/MS acquisition rate for ETD-based methods. Our evaluations of ion-ion reactions conducted at higher Mathieu q-values (relative to the reagent anion at m/z 202) showed that we can increase reaction rate and use shorter time constants to conduct ETD reactions approximately six times faster without a loss of spectral quality. We combined the increase in spectra acquisition rate afforded by these ETD reaction parameter changes with improvements to how the Orbitrap Tribrid MS system parallelizes MS operations and manipulations of various ion populations. These two changes, both implemented on the newest generation of commercially available Tribrid instruments, the Orbitrap Apex, enabled significantly more O-glycopeptide identifications than analogous methods available on the Orbitrap Ascend. Our experiments also showed that spectral acquisition rate is not the only driver for improving glycopeptide identifications. Sufficient precursor ion accumulation time is also needed to collect enough ions for glycopeptide identification. With this requirement for sensitivity, the faster ETD-based methods we developed on the Orbitrap Apex can now compete with sceHCD based methods for N-glycopeptide identifications while sustaining their significant advantage for O-glycopeptide identifications. We note that the balancing of ETD-based acquisition rates are uniquely enabled by the dual ion-routing multipole architecture of the newest Orbitrap Tribrid systems, although the cost-benefit tradeoffs for ETD-based methods will need to be evaluated on systems where necessary ion accumulation times are significantly shorter. Our experiments indicate that these more expedient ETD-based methods can become the universal MS/MS dissociation choice for both N- and O-glycopeptide characterization. Future work will be needed to capitalize on this decision, however. Currently, search algorithms are largely geared to work with HCD-like spectra for glycopeptide identification, even if ETD-based methods are used for localization. New strategies to de-emphasize the reliance on non-modified b/y-type ions for identifications will be needed. Improvement in glycopeptide search algorithms is a well-recognized need and on-going effort^82^, and we hope this study can provide more reasons to build tools that can capitalize on the information-rich spectra ETD-based fragmentation can provide for glycopeptides. Additionally, this work does not address limitations in how to elucidate glycan topology for various glycan compositions at each glycosite, but we envision electron-based dissociation methods will play a key role in future efforts toward this goal for structural glycoproteomics.

## SUPPORTING INFORMATION

Supporting Information is available free of charge online. This information includes Figure S1. Example spectra approximating multiples of tau (τ); Figure S2. O-glycopeptide precursor ion depletion curves at different q-values; Figure S3. Angiotensin precursor ion depletion curves at different q-values; Figure S4. Relationship between reaction rate, q-value, and low mass cutoff. Figure S5. Kinetic dependence on charge state; Figure S6. Considering multiples of the time constant, tau (τ); Figure S7. Behavior of precursor and charge reduced precursor ions under ETD and EThcD40 conditions for triply protontated O-glycopeptide precursor ions; Figure S8. Retention of low m/z oxonium ions; Figure S9. Comparing sceHCD and EThcD methods on the Orbitrap Apex.

## ACKNOWLEDGEMENTS

Research reported in this publication was supported by the U.S. National Science Foundation under CAREER Award 2540459 (N.M.R.) and an ASMS Research Award (N.M.R.). We thank Henrik Clausen, Yoshiki Narimatsu, and Thapakorn Jaroentomeechai for providing the glycocarrier sample, and Jacob Russell for helping with a tryptic digestion of the glycocarrier. The mass spectrometry raw data and search results have been deposited to the ProteomeXchange Consortium via the MassIVE partner repository with the dataset identifier MSV000102759.

## FOR TOC ONLY

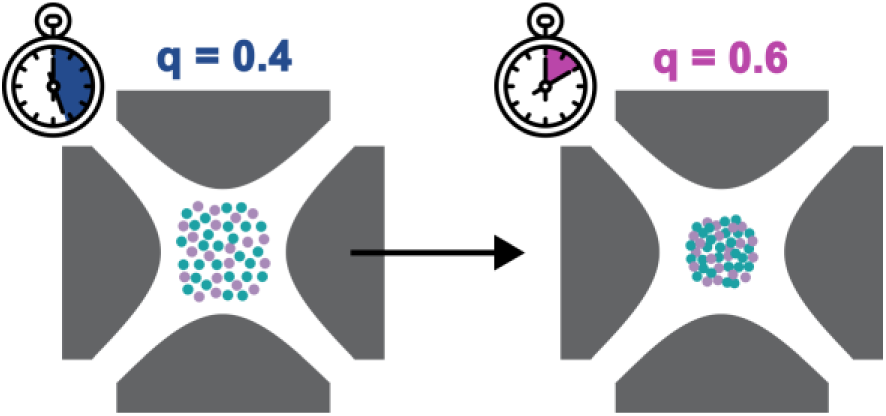

## REFERENCES

(1) Syka, J. E. P.; Coon, J. J.; Schroeder, M. J.; Shabanowitz, J.; Hunt, D. F. Peptide and Protein Sequence Analysis by Electron Transfer Dissociation Mass Spectrometry. Proc. Natl. Acad. Sci. U. S. A. 2004, 101 (26), 9528–9533. 10.1073/pnas.0402700101.

(2) Udeshi, N. D.; Compton, P. D.; Shabanowitz, J.; Hunt, D. F.; Rose, K. L. Methods for Analyzing Peptides and Proteins on a Chromatographic Timescale by Electron-Transfer Dissociation Mass Spectrometry. Nat. Protoc. 2008, 3 (11), 1709–1717. 10.1038/nprot.2008.159.

(3) Lermyte, F.; Valkenborg, D.; Loo, J. A.; Sobott, F. Radical Solutions: Principles and Application of Electron-Based Dissociation in Mass Spectrometry-Based Analysis of Protein Structure. Mass Spectrometry Reviews. Wiley-Blackwell February 9, 2018. 10.1002/mas.21560.

(4) Riley, N. M.; Coon, J. J. The Role of Electron Transfer Dissociation in Modern Proteomics. Anal. Chem. 2018, 90 (1), 40–64. 10.1021/acs.analchem.7b04810.

(5) Coon, J. J. Collisions or Electrons? Protein Sequence Analysis in the 21st Century. Anal. Chem. 2009, 81 (9), 3208–3215. 10.1021/ac802330b.

(6) Myers, S. A.; Daou, S.; Affar, E. B.; Burlingame, A. Electron Transfer Dissociation (ETD): The Mass Spectrometric Breakthrough Essential for O-GlcNAc Protein Site Assignments-a Study of the O-GlcNAcylated Protein Host Cell Factor C1. Proteomics 2013, 13 (6), 982– 991. 10.1002/pmic.201200332.

(7) Doll, S.; Burlingame, A. L. Mass Spectrometry-Based Detection and Assignment of Protein Posttranslational Modifications. ACS Chem. Biol. 2015, 10 (1), 63–71. 10.1021/cb500904b.

(8) Brodbelt, J. S. Deciphering Combinatorial Post-Translational Modifications by Top-down Mass Spectrometry. Curr. Opin. Chem. Biol. 2022, 70, 102180. 10.1016/j.cbpa.2022.102180.

(9) Chapman, J. R. Mass Spectrometry–Based Proteomics for the Masses: Peptide and Protein Identification in the Hunt Laboratory During the 2000’s. Mol. Cell. Proteomics 2024, 23 (12). 10.1016/j.mcpro.2024.100866.

(10) Udeshi, N. D.; Hart, G. W.; Slawson, C. From Fringe to the Mainstream: How ETD MS Brought O-GlcNAc to the *Masses*. Mol. Cell. Proteomics 2024, 23 (11), 100859. 10.1016/j.mcpro.2024.100859.

(11) Coon, J. J.; Marto, J. A.; Syka, J. E. P.; White, F. M. The Hunt Lab Weighs in on Mass Spectrometry–Based Analysis of Protein Posttranslational Modifications. Mol. Cell. Proteomics 2025, 24 (4). 10.1016/j.mcpro.2025.100943.

(12) Wu, F.; Li, W.; Lu, H.; Li, L. Recent Advances in Mass Spectrometry-Based Studies of Post-Translational Modifications in Alzheimer’s Disease. Mol. Cell. Proteomics 2025, 24 (7). 10.1016/j.mcpro.2025.101003.

(13) Poncha, K. F.; Paparella, A. T.; Young, N. L. Unraveling Posttranslational Modification Complexity: Advances in Quantitative Histone Proteoform Mass Spectrometry. Mass Spectrom. Rev. 2026, 45 (4), 769–799. 10.1002/mas.21941.

(14) Jiang, Y.; Rex, D. A. B.; Schuster, D.; Neely, B. A.; Rosano, G. L.; Volkmar, N.; Momenzadeh, A.; Peters-Clarke, T. M.; Egbert, S. B.; Kreimer, S.; Doud, E. H.; Crook, O. M.; Yadav, A. K.; Vanuopadath, M.; Hegeman, A. D.; Mayta, M. L.; Duboff, A. G.; Riley, N. M.; Moritz, R. L.; Meyer, J. G. Comprehensive Overview of Bottom-Up Proteomics Using Mass Spectrometry. ACS Meas. Sci. Au 2024, 4 (4), 338–417. 10.1021/acsmeasuresciau.3c00068.

(15) Reiding, K. R.; Bondt, A.; Franc, V.; Heck, A. J. R. The Benefits of Hybrid Fragmentation Methods for Glycoproteomics. TrAC - Trends Anal. Chem. 2018, *108*, 10.1016/j.trac.2018.09.007. 10.1016/j.trac.2018.09.007.

(16) Ruhaak, L. R.; Xu, G.; Li, Q.; Goonatilleke, E.; Lebrilla, C. B. Mass Spectrometry Approaches to Glycomic and Glycoproteomic Analyses. Chem. Rev. 2018, 118 (17), 7886–7930. 10.1021/acs.chemrev.7b00732.

(17) Bagdonaite, I.; Malaker, S. A.; Polasky, D. A.; Riley, N. M.; Schjoldager, K.; Vakhrushev, S. Y.; Halim, A.; Aoki-Kinoshita, K. F.; Nesvizhskii, A. I.; Bertozzi, C. R.; Wandall, H. H.; Parker, B. L.; Thaysen-Andersen, M.; Scott, N. E. Glycoproteomics. Nat. Rev. Methods Primer 2022, 2 (1), 48. 10.1038/s43586-022-00128-4.

(18) Sandilya, V.; Sahioun, S.; Suma, T. A.; Adewolu, A.; Koraich, A.; Nishe, S. S.; Oji, E.; Gutierrez Reyes, C. D.; Mechref, Y. Advances in Fragmentation Techniques for Glycomics and Glycoproteomics. Mass Spectrom. Rev. 2026, 45 (4), 851–873. 10.1002/mas.70011.

(19) Schjoldager, K. T.; Narimatsu, Y.; Joshi, H. J.; Clausen, H. Global View of Human Protein Glycosylation Pathways and Functions. Nat. Rev. Mol. Cell Biol. 2020 2112 2020, 21 (12), 729–749. 10.1038/s41580-020-00294-x.

(20) Haan, N. de; Nielsen, M. I.; Wandall, H. H. Reading and Writing the Human Glycocode. Annu. Rev. Biochem. 2024, 93 (Volume 93, 2024), 529–564. 10.1146/annurev-biochem-030122-044347.

(21) Fadda, E.; Singh, O.; Schulz, B. L. Heterogeneity of Glycoproteins: Why Does It Matter and How to Account for It. Curr. Opin. Struct. Biol. 2026, 99, 103296. 10.1016/j.sbi.2026.103296.

(22) Chau, T. H.; Chernykh, A.; Kawahara, R.; Thaysen-Andersen, M. Critical Considerations in *N*-Glycoproteomics. Curr. Opin. Chem. Biol. 2023, 73, 102272. 10.1016/j.cbpa.2023.102272.

(23) Darula, Z.; Medzihradszky, K. F. Analysis of Mammalian O-Glycopeptides - We Have Made a Good Start, but There Is a Long Way to Go. Mol. Cell. Proteomics 2018, 17 (1), 2– 17. 10.1074/mcp.MR117.000126.

(24) Riley, N. M.; Malaker, S. A. O-Glycoproteomics: Methods, Challenges, and New Opportunities. 2024. 10.1039/9781839166433-00118.

(25) Chernykh, A.; Kawahara, R.; Thaysen-Andersen, M. Towards Structure-Focused Glycoproteomics. Biochem. Soc. Trans. 2021. 10.1042/bst20200222.

(26) Yu, Q.; Wang, B.; Chen, Z.; Urabe, G.; Glover, M. S.; Shi, X.; Guo, L. W.; Kent, K. C.; Li, L. Electron-Transfer/Higher-Energy Collision Dissociation (EThcD)-Enabled Intact Glycopeptide/Glycoproteome Characterization. J. Am. Soc. Mass Spectrom. 2017, 28 (9), 1751–1764. 10.1007/s13361-017-1701-4.

(27) Glover, M. S.; Yu, Q.; Chen, Z.; Shi, X.; Kent, K. C.; Li, L. Characterization of Intact Sialylated Glycopeptides and Phosphorylated Glycopeptides from IMAC Enriched Samples by EThcD Fragmentation: Toward Combining Phosphoproteomics and Glycoproteomics. Int. J. Mass Spectrom. 2017. 10.1016/J.IJMS.2017.09.002.

(28) Zhang, Y.; Xie, X.; Zhao, X.; Tian, F.; Lv, J.; Ying, W.; Qian, X. Systems Analysis of Singly and Multiply *O*-Glycosylated Peptides in the Human Serum Glycoproteome via EThcD and HCD Mass Spectrometry. J. Proteomics 2018, 170, 14–27. 10.1016/j.jprot.2017.09.014.

(29) Riley, N. M.; Hebert, A. S.; Westphall, M. S.; Coon, J. J. Capturing Site-Specific Heterogeneity with Large-Scale N-Glycoproteome Analysis. Nat. Commun. 2019, 10 (1), 1311. 10.1038/s41467-019-09222-w.

(30) Zhang, R.; Zhu, J.; Lubman, D. M.; Mechref, Y.; Tang, H. GlycoHybridSeq: Automated Identification of N-Linked Glycopeptides Using Electron Transfer/High-Energy Collision Dissociation (EThcD). J. Proteome Res. 2021, 20 (6), 3345–3352. 10.1021/acs.jproteome.1c00245.

(31) Mao, Y.; Wang, S.; Zhao, Y.; Konstantinidi, A.; Sun, L.; Ye, Z.; Vakhrushev, S. Y. Systematic Evaluation of Fragmentation Methods for Unlabeled and Isobaric Mass Tag-Labeled O-Glycopeptides. Anal. Chem. 2021, 93 (32), 11167–11175. 10.1021/acs.analchem.1c01696.

(32) Downs, M.; Curran, J.; Zaia, J.; Sethi, M. K. Analysis of Complex Proteoglycans Using Serial Proteolysis and EThcD Provides Deep N- and O-Glycoproteomic Coverage. Anal. Bioanal. Chem. 2023, 415 (28), 6995–7009. 10.1007/s00216-023-04934-x.

(33) Darula, Z.; Pap, Á.; Medzihradszky, K. F. Extended Sialylated O-Glycan Repertoire of Human Urinary Glycoproteins Discovered and Characterized Using Electron-Transfer/Higher-Energy Collision Dissociation. J. Proteome Res. 2019, 18 (1), 280–291. 10.1021/acs.jproteome.8b00587.

(34) Kuo, C.-W.; Guu, S.-Y.; Khoo, K.-H. Distinctive and Complementary MS2 Fragmentation Characteristics for Identification of Sulfated Sialylated N-Glycopeptides by nanoLC-MS/MS Workflow. J. Am. Soc. Mass Spectrom. 2018, 29 (6), 1166–1178. 10.1007/s13361-018-1919-9.

(35) Riley, N. M.; Malaker, S. A.; Driessen, M.; Bertozzi, C. R. Optimal Dissociation Methods Differ for N-and O-Glycopeptides. J. Proteome Res. 2020. 10.1021/acs.jproteome.0c00218.

(36) Pap, A.; Klement, E.; Hunyadi-Gulyas, E.; Darula, Z.; Medzihradszky, K. F. Status Report on the High-Throughput Characterization of Complex Intact O-Glycopeptide Mixtures. J. Am. Soc. Mass Spectrom. 2018, 29 (6), 1210–1220. 10.1007/s13361-018-1945-7.

(37) Chalkley, R. J.; Medzihradszky, K. F.; Darula, Z.; Pap, A.; Baker, P. R. The Effectiveness of Filtering Glycopeptide Peak List Files for Y Ions. *Mol*. Omics 2020, 16 (2), 147–155. 10.1039/c9mo00178f.

(38) Riley, N. M.; Malaker, S. A.; Bertozzi, C. R. Electron-Based Dissociation Is Needed for O-Glycopeptides Derived from OpeRATOR Proteolysis. Anal. Chem. 2020, 92 (22), 14878– 14884. 10.1021/ACS.ANALCHEM.0C02950.

(39) Pap, A.; Kiraly, I. E.; Medzihradszky, K. F.; Darula, Z. Multiple Layers of Complexity in O-Glycosylation Illustrated With the Urinary Glycoproteome. Mol. Cell. Proteomics 2022, 21 (12). 10.1016/j.mcpro.2022.100439.

(40) Chongsaritsinsuk, J.; Rangel-Angarita, V.; Lucas, T. M.; Mahoney, K. E.; Enny, O. M.; Katemauswa, M.; Malaker, S. A. Quantification and Site-Specific Analysis of Co-Occupied N- and O-Glycopeptides. J. Proteome Res. 2024, 23 (12), 5449–5461. 10.1021/acs.jproteome.4c00574.

(41) Peters-Clarke, T. M.; Coon, J. J.; Riley, N. M. Instrumentation at the Leading Edge of Proteomics. Anal. Chem. 2024, 96 (20), 7976–8010. 10.1021/acs.analchem.3c04497.

(42) Earley, L.; Anderson, L. C.; Bai, D. L.; Mullen, C.; Syka, J. E. P.; English, A. M.; Dunyach, J.-J.; Stafford, G. C.; Shabanowitz, J.; Hunt, D. F.; Compton, P. D. Front-End Electron Transfer Dissociation: A New Ionization Source. Anal. Chem. 2013, 85 (17), 8385–8390. 10.1021/ac401783f.

(43) Downs, M.; Xia, C.; Lin, C.; Zaia, J. Optimization of LC-ExD Methods for Glycopeptides on an Omnitrap-Orbitrap Platform. Anal. Bioanal. Chem. 2026, 418 (9), 2721–2729. 10.1007/s00216-026-06400-w.

(44) Levin, N.; Polasky, D. A.; Li, K.; Nesvizhskii, A. I.; Mohammed, S. Evaluating the Performance of Photon- and Electron-Based Fragmentation Methods in Omnitrap-LCMS Analysis of N-Glycopeptides. Anal. Chem. 2026, 98 (27), 20043–20054. 10.1021/acs.analchem.6c00435.

(45) Li, R.; Xia, C.; Wu, S.; Downs, M. J.; Tong, H.; Tursumamat, N.; Zaia, J.; Costello, C. E.; Lin, C.; Wei, J. Direct and Detailed Site-Specific Glycopeptide Characterization by Higher-Energy Electron-Activated Dissociation Tandem Mass Spectrometry. Anal. Chem. 2024, 96 (3), 1251–1258. 10.1021/acs.analchem.3c04484.

(46) Searfoss, R. M.; Zahn, E.; Lin, Z.; Garcia, B. A. Establishing a Top-Down Proteomics Platform on a Time-of-Flight Instrument with Electron-Activated Dissociation. J. Proteome Res. 2025, 24 (3), 1230–1240. 10.1021/acs.jproteome.4c00874.

(47) Manly, L. S.; Roberts, A. M.; Beckman, J. S.; Roberts, B. R. Electron Capture Dissociation for Discovery Top-Down Proteomics of Peptides and Small Proteins on Chromatographic Time Scales. J. Am. Soc. Mass Spectrom. 2025, 36 (10), 2079–2093. 10.1021/jasms.5c00116.

(48) Beckman, J. S.; Voinov, V. G.; Hare, M.; Sturgeon, D.; Vasil’ev, Y.; Oppenheimer, D.; Shaw, J. B.; Wu, S.; Glaskin, R.; Klein, C.; Schwarzer, C.; Stafford, G. Improved Protein and PTM Characterization with a Practical Electron-Based Fragmentation on Q-TOF Instruments. J. Am. Soc. Mass Spectrom. 2021, 32 (8), 2081–2091. 10.1021/JASMS.0C00482.

(49) Ollivier, S.; Schuster, D.; van Rijswijck, D. M. H.; Stroganova, I.; Smyrnakis, A.; Fiala, J.; Kosmopoulou, M.; Suckau, D.; Pengelley, S.; Raether, O.; Greisch, J.-F.; Papanastasiou, D.; Heck, A. J. R. A Prototype timsOmni Platform Enables Confident Annotation of the Key Hypervariable CDR3 Regions of IgG Immunoglobulins Using Low- and High-Energy Electron-Based Fragmentation. Anal. Chem. 2026, 98 (15), 11070–11082. 10.1021/acs.analchem.5c07288.

(50) Baba, T.; Zhang, Z.; Liu, S.; Burton, L.; Ryumin, P.; Le Blanc, J. C. Y. Localization of Multiple O-Linked Glycans Exhibited in Isomeric Glycopeptides by Hot Electron Capture Dissociation. J. Proteome Res. 2022, 21 (10), 2462–2471. 10.1021/acs.jproteome.2c00378.

(51) Macauslane, K. L.; Pegg, C. L.; Nouwens, A. S.; Kerr, E. D.; Seitanidou, J.; Schulz, B. L. Electron-Activated Dissociation and Collision-Induced Dissociation Glycopeptide Fragmentation for Improved Glycoproteomics. Anal. Chem. 2024, 96 (27), 10986–10994. 10.1021/acs.analchem.4c01450.

(52) Veth, T. S.; Sutherland, E.; Markuson, K. A.; Zhang, R.; Duboff, A. G.; Huang, J.; Bergen, D.; Lee, A. E.; Melani, R. D.; Canterbury, J. D.; Zabrouskov, V.; McAlister, G. C.; Mullen, C.; Riley, N. M. Improvements in Glycoproteomics through Architecture Changes to the Orbitrap Tribrid MS Platform. Anal. Chem. 2025, 97 (22), 11413–11423. 10.1021/acs.analchem.4c06370.

(53) Kessler, A. L.; Fort, K. L.; Resemann, H. C.; Krüger, P.; Wang, C.; Koch, H.; Hauschild, J.-P.; Marino, F.; Heck, A. J. R. Increased EThcD Efficiency on the Hybrid Orbitrap Excedion Pro Mass Analyzer Extends the Depth in Identification and Sequence Coverage of HLA Class I Immunopeptidomes. Mol. Cell. Proteomics 2025, 24 (9), 101049. 10.1016/j.mcpro.2025.101049.

(54) Veth, T. S.; Riley, N. M. Accelerating the Stride toward Functional Glycoproteomics. Trends Biochem. Sci. 2025, 50 (7), 550–552. 10.1016/j.tibs.2025.04.002.

(55) Sun, Z.; Lin, D. C.; Hu, Y.; Zhang, H. From Identification to Quantification: A Step-by-Step Workflow for Intact Glycopeptide Analysis by Mass Spectrometry. Anal. Chim. Acta 2026, 1383, 344909. 10.1016/j.aca.2025.344909.

(56) Singh, C.; Zampronio, C. G.; Creese, A. J.; Cooper, H. J. Higher Energy Collision Dissociation (HCD) Product Ion-Triggered Electron Transfer Dissociation (ETD) Mass Spectrometry for the Analysis of N-Linked Glycoproteins. J. Proteome Res. 2012, 11 (9), 4517–4525. 10.1021/pr300257c.

(57) Zhao, P.; Viner, R.; Teo, C. F.; Boons, G. J.; Horn, D.; Wells, L. Combining High-Energy C-Trap Dissociation and Electron Transfer Dissociation for Protein O-GlcNAc Modification Site Assignment. J. Proteome Res. 2011, 10 (9), 4088–4104. 10.1021/pr2002726.

(58) Wu, S.-W.; Pu, T.-H.; Viner, R.; Khoo, K.-H. Novel LC-MS ^2^ Product Dependent Parallel Data Acquisition Function and Data Analysis Workflow for Sequencing and Identification of Intact Glycopeptides. Anal. Chem. 2014, 86 (11), 5478–5486. 10.1021/ac500945m.

(59) Saba, J.; Dutta, S.; Hemenway, E.; Viner, R. Increasing the Productivity of Glycopeptides Analysis by Using Higher-Energy Collision Dissociation-Accurate Mass-Product-Dependent Electron Transfer Dissociation. Int. J. Proteomics 2012, 2012, 1–7. 10.1155/2012/560391.

(60) Zhu, J.; Chen, Z.; Zhang, J.; An, M.; Wu, J.; Yu, Q.; Skilton, St. J.; Bern, M.; Ilker Sen, K.; Li, L.; Lubman, D. M. Differential Quantitative Determination of Site-Specific Intact N-Glycopeptides in Serum Haptoglobin between Hepatocellular Carcinoma and Cirrhosis Using LC-EThcD-MS/MS. J. Proteome Res. 2019, 18 (1), 359–371.

(61) Cao, Q.; Yu, Q.; Liu, Y.; Chen, Z.; Li, L. Signature-Ion-Triggered Mass Spectrometry Approach Enabled Discovery of N- and O-Linked Glycosylated Neuropeptides in the Crustacean Nervous System. J. Proteome Res. 2020, 19 (2), 634–643. 10.1021/acs.jproteome.9b00525.

(62) Sutherland, E.; Veth, T. S.; Barshop, W. D.; Russell, J. H.; Kothlow, K.; Canterbury, J. D.; Mullen, C.; Bergen, D.; Huang, J.; Zabrouskov, V.; Huguet, R.; McAlister, G. C.; Riley, N. M. Autonomous Dissociation-Type Selection for Glycoproteomics Using a Real-Time Library Search. J. Proteome Res. 2024, 23 (12), 5606–5614. 10.1021/acs.jproteome.4c00723.

(63) Jaroentomeechai, T.; Karlsson, R.; Goerdeler, F.; Teoh, F. K. Y.; Grønset, M. N.; de Wit, D.; Chen, Y.-H.; Furukawa, S.; Psomiadou, V.; Hurtado-Guerrero, R.; Vidal-Calvo, E. E.; Salanti, A.; Boltje, T. J.; van den Bos, L. J.; Wunder, C.; Johannes, L.; Schjoldager, K. T.; Joshi, H. J.; Miller, R. L.; Clausen, H.; Vakhrushev, S. Y.; Narimatsu, Y. Mammalian Cell-Based Production of Glycans, Glycopeptides and Glycomodules. Nat. Commun. 2024, 15 (1), 9668. 10.1038/s41467-024-53738-9.

(64) Veth, T. S.; Kothlow, K.; Riley, N. M. Understanding m/z Range Settings for MS/MS Scans: A Case Study with Intact Glycopeptides. J. Am. Soc. Mass Spectrom. 2026, 37 (4), 872– 883. 10.1021/jasms.5c00352.

(65) Hsu, C.; Heil, L. R.; Wen, B.; McAlister, G.; Merrihew, G.; Remes, P. M.; Plubell, D. L.; Melani, R.; Zabrouskov, V.; MacCoss, M. J. Revisiting Resonance-Excitation Collision-Induced Dissociation for Data-Independent Acquisition. bioRxiv 2026, 2026.05.31.729078. 10.64898/2026.05.31.729078.

(66) Lu, L.; Riley, N. M.; Shortreed, M. R.; Bertozzi, C. R.; Smith, L. M. O-Pair Search with MetaMorpheus for O-Glycopeptide Characterization. Nat. Methods 2020, 17 (11), 1133– 1138. 10.1038/s41592-020-00985-5.

(67) Riley, N.; Bertozzi, C. Deciphering O-Glycoprotease Substrate Preferences with O-Pair Search. *Mol*. Omics 2022, 10.1039/D2MO00244B. 10.1039/D2MO00244B.

(68) Polasky, D. A.; Yu, F.; Teo, G. C.; Nesvizhskii, A. I. Fast and Comprehensive N- and O-Glycoproteomics Analysis with MSFragger-Glyco. Nat. Methods 2020, 17 (11), 1125–1132. 10.1038/s41592-020-0967-9.

(69) Kothlow, K.; Schramm, H. M.; Markuson, K. A.; Russell, J. H.; Sutherland, E.; Veth, T. S.; Zhang, R.; Duboff, A. G.; Tejus, V. R.; McDermott, L. E.; Dräger, L. S.; Riley, N. M. Extracting Informative Glycan-Specific Ions From Glycopeptide MS/MS Spectra With GlyCounter. Mol. Cell. Proteomics 2025, 24 (12), 101085. 10.1016/j.mcpro.2025.101085.

(70) McLuckey, S. A.; Stephenson, J. L.; Asano, K. G. Ion/Ion Proton-Transfer Kinetics: Implications for Analysis of Ions Derived from Electrospray of Protein Mixtures. Anal. Chem. 1998, 70 (6), 1198–1202.

(71) Xia, Y.; Wu, J.; Mcluckey, S. A.; Londry, F. A.; Hager, J. W. Mutual Storage Mode Ion/Ion Reactions in a Hybrid Linear Ion Trap. J. Am. Soc. Mass Spectrom. 2005, 16 (1), 71–81. 10.1016/j.jasms.2004.09.017.

(72) McLuckey, S. A.; Huang, T.-Y. Ion/Ion Reactions: New Chemistry for Analytical MS. Anal. Chem. 2009, 81 (21), 8669–8676. 10.1021/ac9014935.

(73) Stephenson, J. L.; McLuckey, S. A. Ion/Ion Reactions in the Gas Phase: Proton Transfer Reactions Involving Multiply-Charged Proteins. J. Am. Chem. Soc. 1996, 118 (31), 7390– 7397. 10.1021/ja9611755.

(74) Rose, C. M.; Rush, M. J. P.; Riley, N. M.; Merrill, A. E.; Kwiecien, N. W.; Holden, D. D.; Mullen, C.; Westphall, M. S.; Coon, J. J. A Calibration Routine for Efficient ETD in Large-Scale Proteomics. J. Am. Soc. Mass Spectrom. 2015, 26 (11), 1848–1857. 10.1007/s13361-015-1183-1.

(75) Kelstrup, C. D.; Frese, C.; Heck, A. J. R.; Olsen, J. V.; Nielsen, M. L. Analytical Utility of Mass Spectral Binning in Proteomic Experiments by SPectral Immonium Ion Detection (SPIID)*. Mol. Cell. Proteomics 2014, 13 (8), 1914–1924. 10.1074/mcp.O113.035915.

(76) Couttas, T. A.; Raftery, M. J.; Bernardini, G.; Wilkins, M. R. Immonium Ion Scanning for the Discovery of Post-Translational Modifications and Its Application to Histones. J. Proteome Res. 2008, 7 (7), 2632–2641. 10.1021/pr700644t.

(77) Compton, P. D.; Strukl, J. V.; Bai, D. L.; Shabanowitz, J.; Hunt, D. F. Optimization of Electron Transfer Dissociation via Informed Selection of Reagents and Operating Parameters. Anal. Chem. 2012, 84 (3), 1781–1785. 10.1021/ac202807h.

(78) Vainauskas, S.; Guntz, H.; McLeod, E.; McClung, C.; Ruse, C.; Shi, X.; Taron, C. H. A Broad-Specificity O-Glycoprotease That Enables Improved Analysis of Glycoproteins and Glycopeptides Containing Intact Complex O-Glycans. Anal. Chem. 2022, 94 (2), 1060– 1069. 10.1021/acs.analchem.1c04055.

(79) Chan, T.-W. D.; Ip, W. H. H. Optimization of Experimental Parameters for Electron Capture Dissociation of Peptides in a Fourier Transform Mass Spectrometer. J. Am. Soc. Mass Spectrom. 2002, 13 (12), 1396–1406. 10.1016/S1044-0305(02)00703-1.

(80) Lu, L.; Riley, N. M.; Shortreed, M. R.; Bertozzi, C. R.; Smith, L. M. O-Pair Search with MetaMorpheus for O-Glycopeptide Characterization. Nat. Methods 2020, 17 (11), 1133– 1138. 10.1038/s41592-020-00985-5.

(81) Polasky, D. A.; Lu, L.; Yu, F.; Li, K.; Shortreed, M. R.; Smith, L. M.; Nesvizhskii, A. I. Quantitative Proteome-Wide O-Glycoproteomics Analysis with FragPipe. Anal. Bioanal. Chem. 2025, 417 (5), 921–930. 10.1007/s00216-024-05382-x.

(82) Kawahara, R.; Chernykh, A.; Alagesan, K.; Bern, M.; Cao, W.; Chalkley, R. J.; Cheng, K.; Choo, M. S.; Edwards, N.; Goldman, R.; Hoffmann, M.; Hu, Y.; Huang, Y.; Kim, J. Y.; Kletter, D.; Liquet, B.; Liu, M.; Mechref, Y.; Meng, B.; Neelamegham, S.; Nguyen-Khuong, T.; Nilsson, J.; Pap, A.; Park, G. W.; Parker, B. L.; Pegg, C. L.; Penninger, J. M.; Phung, T. K.; Pioch, M.; Rapp, E.; Sakalli, E.; Sanda, M.; Schulz, B. L.; Scott, N. E.; Sofronov, G.; Stadlmann, J.; Vakhrushev, S. Y.; Woo, C. M.; Wu, H.-Y.; Yang, P.; Ying, W.; Zhang, H.; Zhang, Y.; Zhao, J.; Zaia, J.; Haslam, S. M.; Palmisano, G.; Yoo, J. S.; Larson, G.; Khoo, K.-H.; Medzihradszky, K. F.; Kolarich, D.; Packer, N. H.; Thaysen-Andersen, M. Community Evaluation of Glycoproteomics Informatics Solutions Reveals High-Performance Search Strategies for Serum Glycopeptide Analysis. Nat. Methods 2021 2021, 18 (11), 1304–1316. 10.1038/s41592-021-01309-x.

