## Supplemental Information for "Unifying N- and O-glycopeptide dissociation methods with expedient electron transfer dissociation"

The mass spectrometry raw data and search results have been deposited to the ProteomeXchange Consortium via the MassIVE partner repository with the dataset identifier MSV000102759.

### Table of Contents

**Figure S1.** Example spectra approximating multiples of tau (τ).
**Figure S2**. O-glycopeptide precursor ion depletion curves at different q-values.
**Figure S3**. Angiotensin precursor ion depletion curves at different q-values.
**Figure S4**. Relationship between reaction rate, q-value, and low mass cutoff.
**Figure S5**. Kinetic dependence on charge state
**Figure S6**. Considering multiples of the time constant, tau (τ).
**Figure S7**. Behavior of precursor and charge reduced precursor ions under ETD and EThcD40 conditions for triply protontated O-glycopeptide precursor ions.
**Figure S8**. Retention of low m/z oxonium ions.
**Figure S9**. Comparing sceHCD and EThcD methods on the Orbitrap Apex.

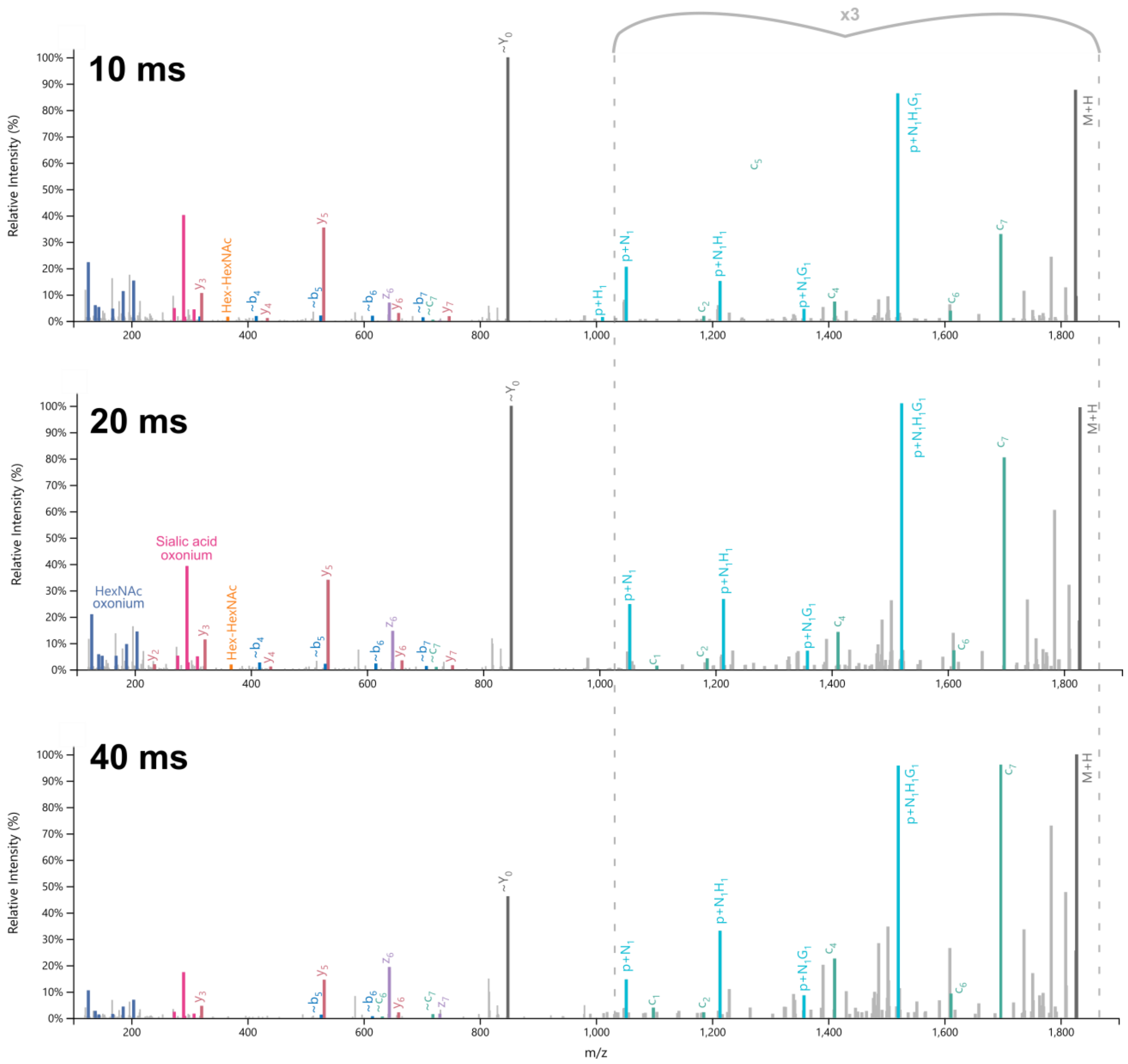


**Figure S1: Example spectra approximating multiples of tau (τ).** Above are EThcD spectra of the doubly charged glycopeptide T[HexNAc(1)Hex(1)NeuGc(2)]SEPLSSK at three different ETD reaction times using a q-value of 0.6. One multiple of the time constant, τ, is 17 ms under these reaction conditions, so these spectra approximate ~0.5τ, 1τ, and 2τ reaction times. Boosts in several of the glycan retained fragments (i.e. c7 and c4) at 20 ms are further realized at 40 ms, but other ions are lost such as the oxonium ions. This tradeoff illustrates a longer-than-needed reaction time that results in an “overreacted precursor”.

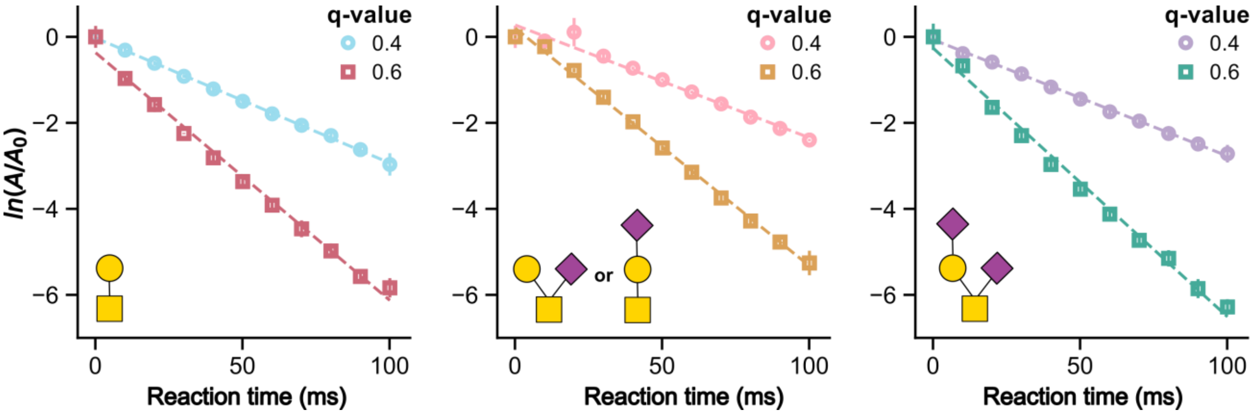


**Figure S2. O-glycopeptide precursor ion depletion curves at different q-values.** These curves show the ratio of remaining precursor ion signal [A / $A_0$] as a function of reaction time in milliseconds. The three graphs show precursor depletion at a q-value of 0.4 and 0.6 for three different doubly charged glycoforms of AEAAAT*PAPAK, where the asterisks deontes the glycan composition shown for the graph. The ETD rate constant (k) is the negative value of the slope.

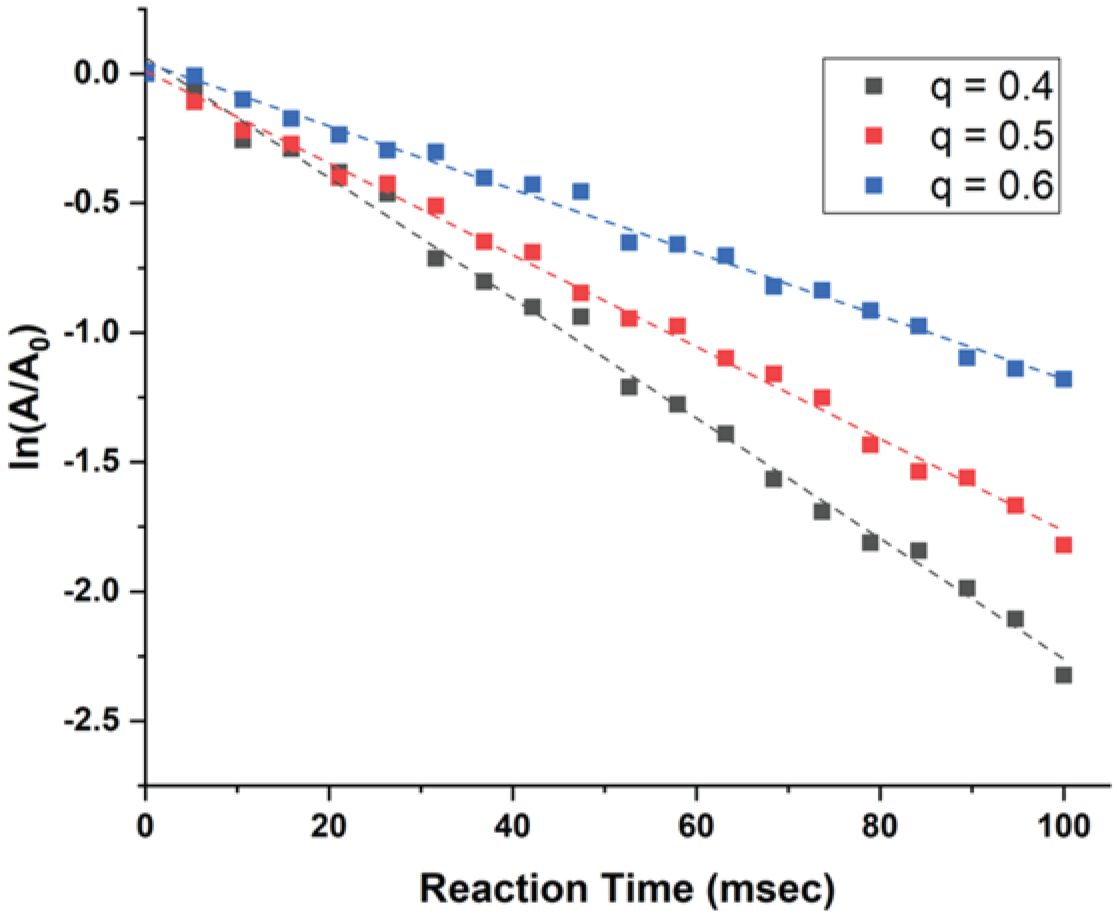


**Figure S3. Angiotensin precursor ion depletion curves at different q-values.** These curves show the ratio of remaining precursor ion signal [A / $A_0$] as a function of reaction time in milliseconds. The graph shows precursor depletion at varying q-values for the triply charged precursor of angiotensin. The ETD rate constant (k) is the negative value of the slope.

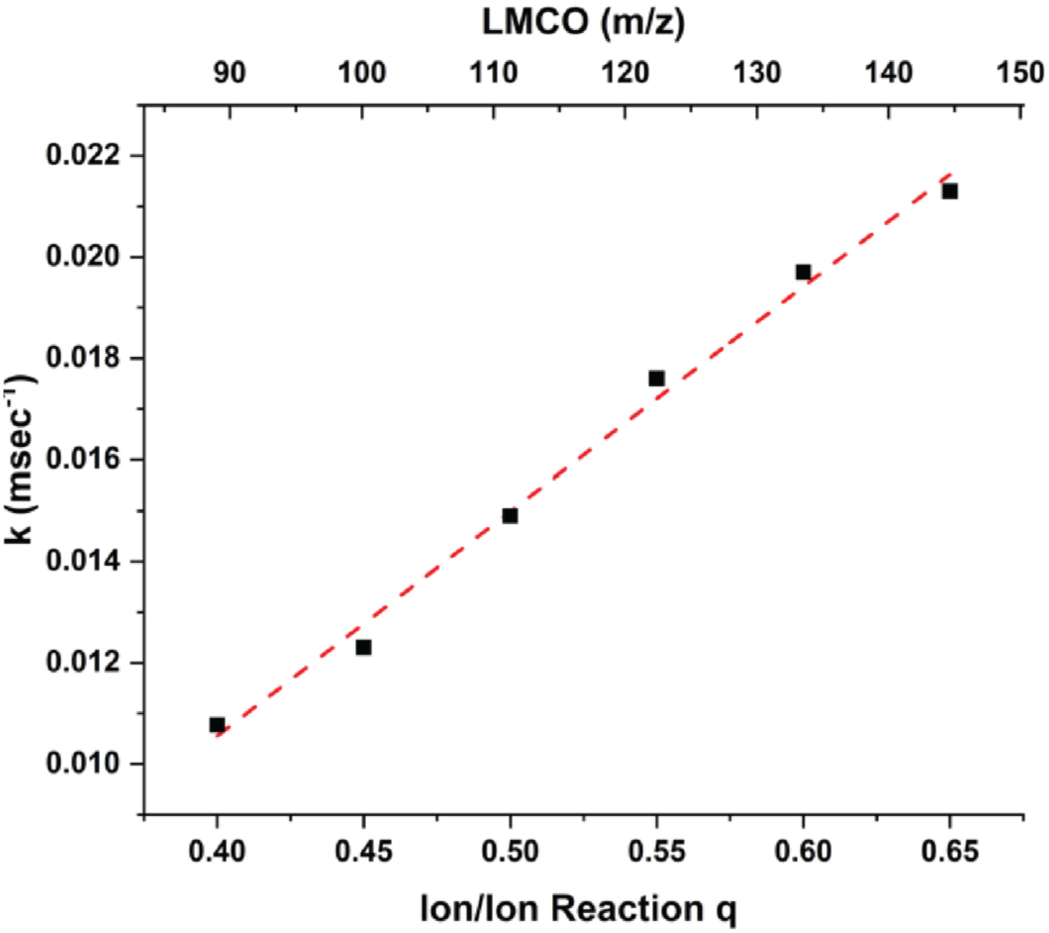


**Figure S4. Relationship between reaction rate, q-value, and low mass cutoff.** This graph shows the relationship between the q-value used during the ion-ion reaction and ETD rate constant, k'. It also shows how the low mass cutoff (LMCO) increases as the q-value increases. These data were generated using the triply protonated precursor cation of angiotensin.

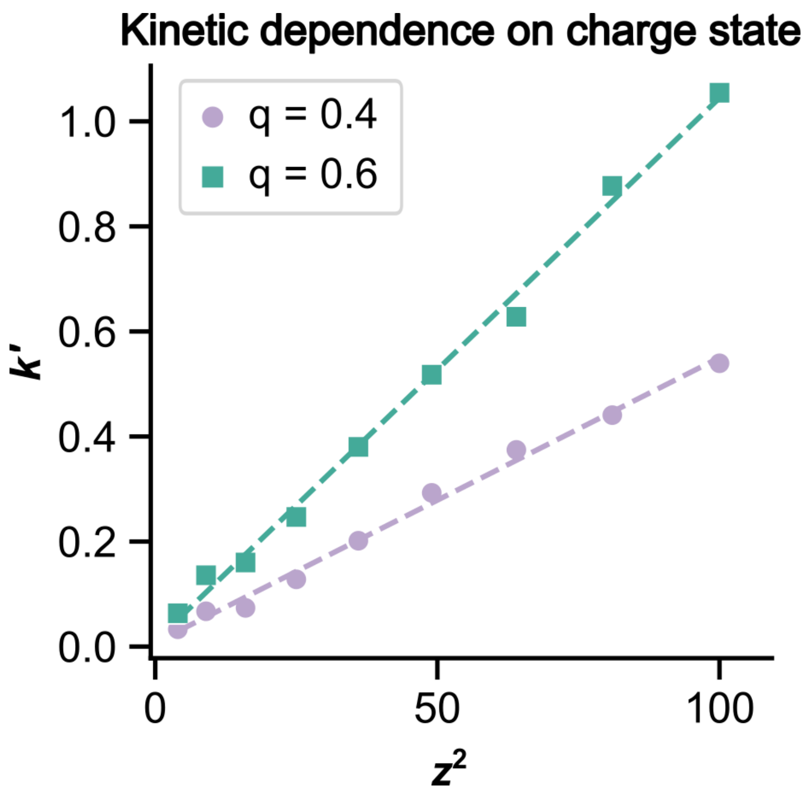


**Figure S5. Kinetic dependence on charge state.** This graph shows the how the ETD reaction rate constant k' scales with charge state in the expected charge-squared relationship. These data were generated using the precursor cations of angiotensin and ubiquitin.

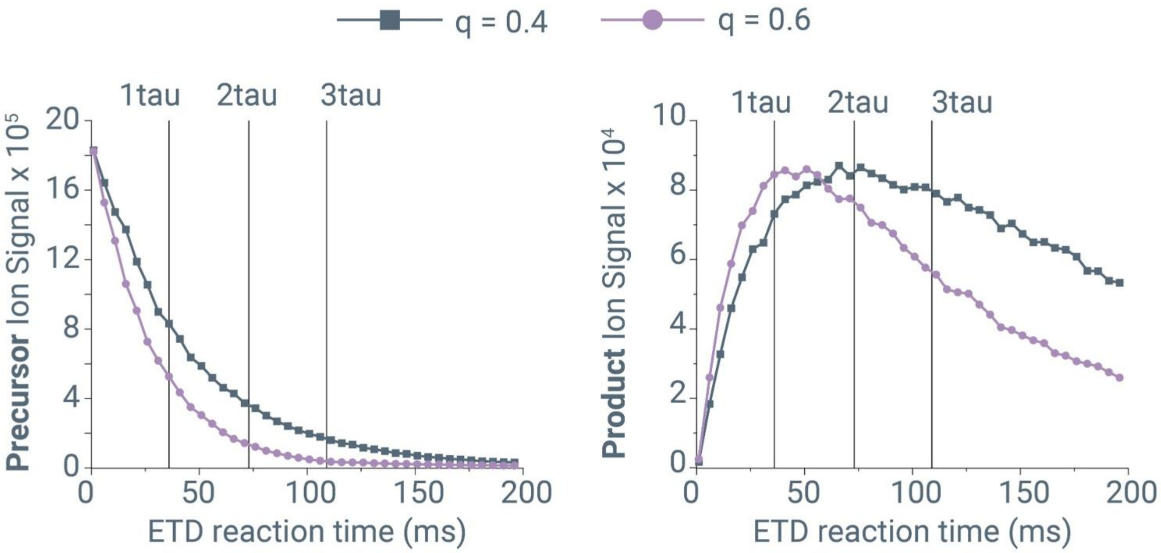


**Figure S6. Considering multiples of the time constant, tau (τ).** Precursor ion depletion (left) and product ion generation (right) as a function of reation time for the triply protonated precursor cation of angiotensin for ion-ion reactoin conducted with a q-value of either 0.4 (gray) or 0.6 (purple). Vertical lines show multiples of the time constant, tau (τ) for the standard ETD reaction conditions (q = 0.4).

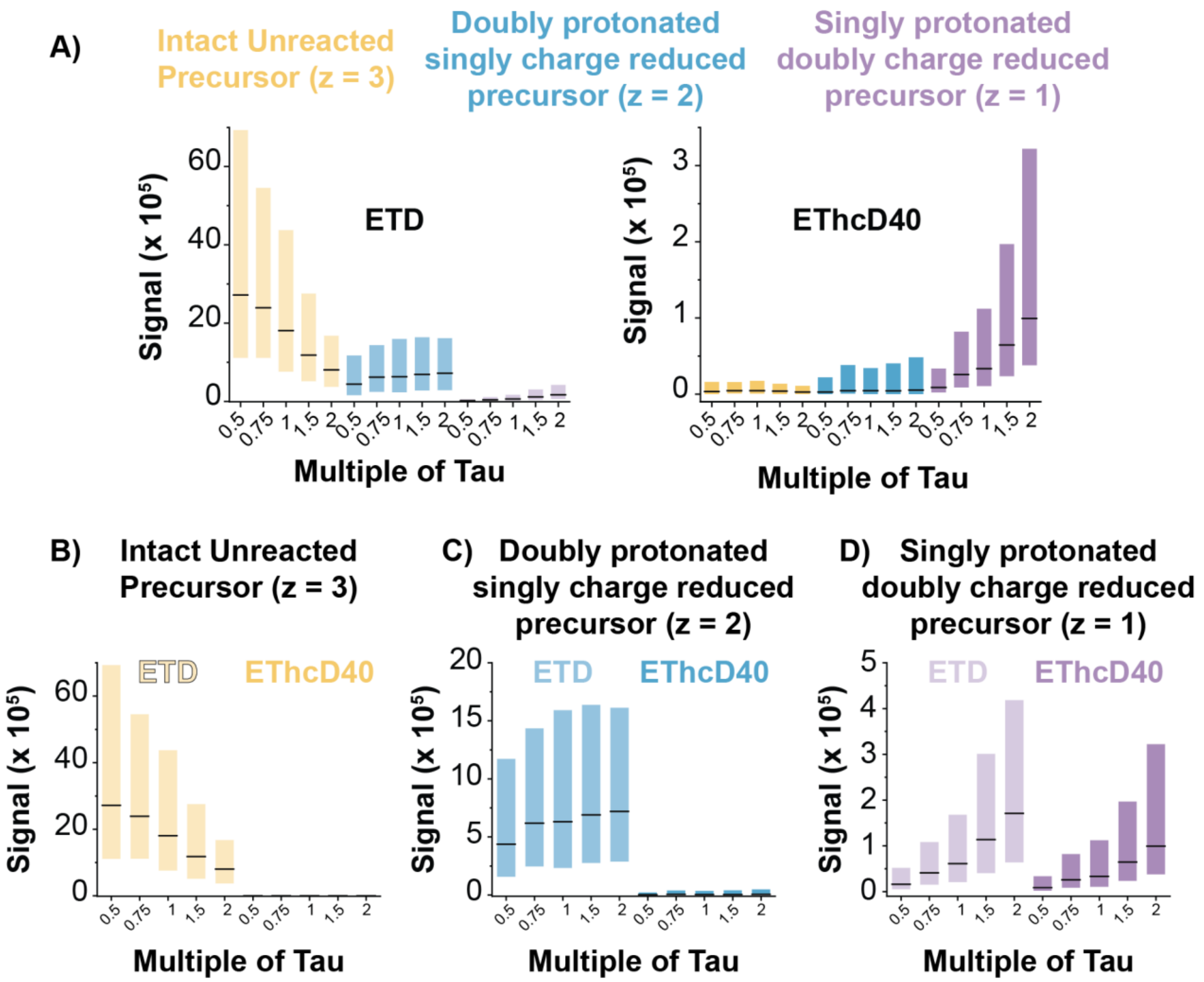


**Figure S7. Behavior of precursor and charge reduced precursor ions under ETD and EThcD40 conditions for triply protontated O-glycopeptide precursor ions. A)** Signal for intact z = 3 unreacted precursor ions (yellow), doubly-charged, singly-charge-reduced precursor ions (z = 2, blue), and singly-charged, doubly-charge-reduced precursor ions (z = 1, purple) are shown for various multiples of tau with q-value = 0.4 for ETD and EThcD40. For ETD, creation and depletion of various ion populations follow expected trends, while in EThcD40, the multiple charged species are depleted more than the singly-charged, doubly-charge-reduced precursor ions (z = 1) during the supplemental HCD activation. Panels **B)**, **C)**, and **D)** show ETD (left five boxes) and EThcD (right five boxes) data on the same graph for each of the three respective populations. The same trends were observed for a reaction q-value = 0.6 (data not shown).

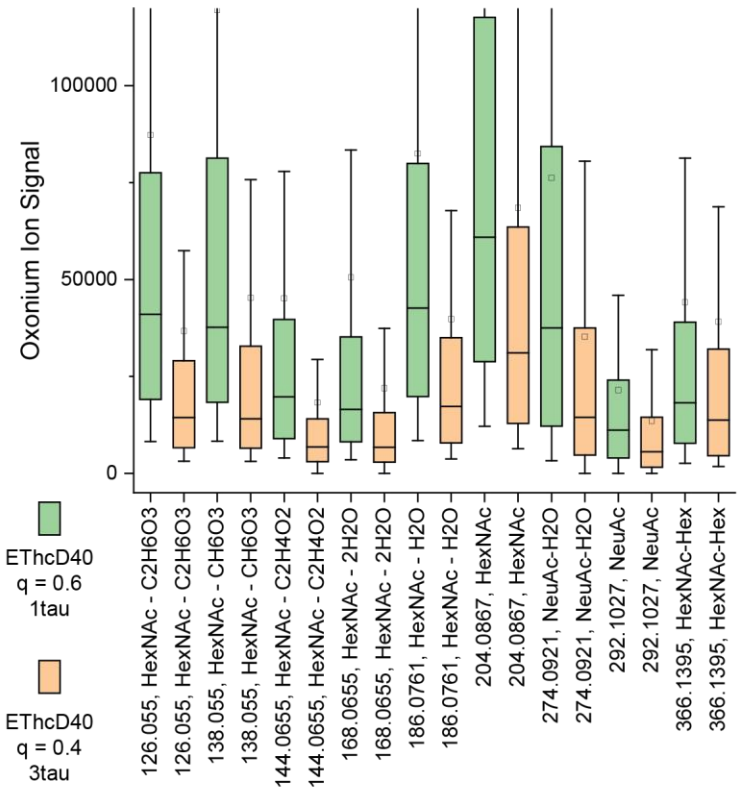


**Figure S8. Retention of low m/z oxonium ions.** Box plots show intensities of low m/z oxonium ions common to glycopeptide spectra for EThcD40 spectra using either a q-value of 0.6 and a single multiple of tau for reaction time (green, new method introduced in this work, n = 9,620 spectra) or a q-value of 0.4 and three multiples of tau for reaction time (orange, standard ETD paramters on commerical Orbitrap systems, n = 6,307 spectra). The slighly higher oxonium ion signal in the q = 0.6 condition is likely due to their being slightly more precursor remaining in the q = 0.6 condition from which to generate oxonium ions during the supplemental HCD step. Regardless, this data shows that retaining low m/z ions in EThcD spectra is not a challenge when increasing the reaction q-value to 0.6.

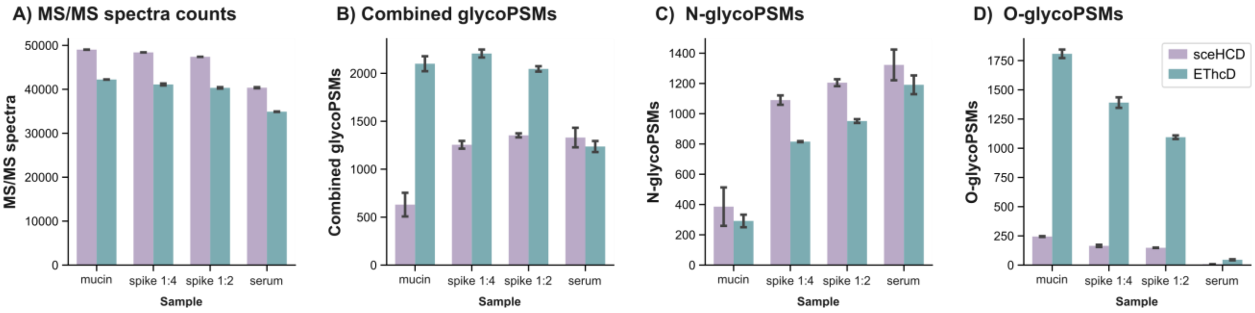


**Figure S9. Comparing sceHCD and EThcD methods on the Orbitrap Apex.** Bar graphs showing the number of **A)** MS/MS spectra, **B)** combined glycopeptide PSMs, **C)** O-glycopeptide PSMs, and **D)** N-glycopeptide PSMs from DDA methods on the Orbitrap Apex using either sceHCD or EThcD40 for MS/MS dissociation. The samples analyzed were either glycopeptides enriched from human serum, a digest of recombinant mucin glycoproteins, or a combination of both to create a complex mixture of both N- and O-glycopeptides. The sceHCD results are shown in purple and EThcD in green. Data were collected in duplicate, and bars show the average of the two replicates, whose values are indicated by the error bars.
